# Virus-host protein-protein interactions between human papillomavirus 16 E6 A1 and D2/D3 sub-lineages: variances and similarities

**DOI:** 10.1101/2020.06.26.169458

**Authors:** Guillem Dayer, Mehran L. Masoom, Melissa Togtema, Ingeborg Zehbe

**Affiliations:** Lakehead University, Biology Department, Thunder Bay, Ontario, Canada; Thunder Bay Regional Health Research Institute, Probe Development and Biomarker Exploration, Thunder Bay, Ontario, Canada; Northern Ontario School of Medicine, Lakehead University, Thunder Bay, Ontario, Canada

## Abstract

High-risk strains of human papillomavirus are causative agents for cervical and other mucosal cancers with type 16 being the most frequent. Compared to the European Prototype (A1, denoted “EP”), the Asian-American (D2/D3, denoted “AA”) sub-lineage or “variant” is reported to have increased abilities to promote carcinogenesis. Few global interactome studies have looked at protein-protein interactions (PPIs) between host proteins and variants of the key transforming E6 protein. We applied a primary human foreskin keratinocyte model transduced with EP and AA variant E6 genes and co-immunoprecipitated expressed E6 proteins along with interacting cellular proteins to detect virus-host binding partners. We reasoned that, due to single nucleotide polymorphisms, AAE6 and EPE6 may have unique PPIs with host cellular proteins—conferring gain or loss of function—resulting in varied abilities to promote carcinogenesis. Using liquid chromatography-mass spectrometry and stringent interactor selection criteria based on the number of peptides, we identified 25 candidates: 6 unique to each of AAE6 and EPE6, along with 13 E6 targets common to both AAE6 and EPE6. We also applied a more inclusive process based on pathway selection and discovered 171 target proteins: 90 unique AAE6 and 61 unique EPE6 along with 20 common E6 targets between the two sub-lineages. Interpretations for both approaches were made using databases such as UniProt, BioGRID and Reactome. Detected E6 targets are implicated in important hallmarks of cancer: deregulating Notch and other signaling, energetics and hypoxia, DNA replication and repair, and immune response. Validation experiments, such as reverse co-immunoprecipitation and RNA interference, are required to substantiate these findings. Here, we provide an unprecedented resource for new research questions in HR HPV biology. The current data also underline our lab’s driving hypothesis that E6, being a “master regulator” in HPV-positive cancers, is an excellent candidate for anti-cancer treatment strategies.

**Author Summary:** Chronic infection with high-risk human papillomavirus (HPV) type 16 is the most prevalent cause of cervical and other mucosal cancers. The E6 oncoproteins of the European Prototype (EP) and the Asian-American (AA) HPV variants differentially promote carcinogenesis. We looked at protein-protein interactions between host proteins and two key HPV variant E6 proteins of these strains to reveal how high risk HPVs cause cancer, based on the proteins they bind to in infected cells. Our methodology combined molecular biology and data mining techniques using widely available databases. We confirmed and discovered novel virus-host associations that explained how HPV AA and EP variants differ in their carcinogenic capabilities, and confirmed the candidacy of the E6 protein as a viable target for HPV therapies.

## Introduction

Human papillomaviruses (HPVs) are double-stranded DNA viruses that infect keratinocytes of skin and mucosal tissues (zur Hausen 2002). Persistent infection with any of the high-risk (HR) types is implicated in almost every case of cervical cancer (Walboomers et al. 1999, Crow 2012), as well as having an aetiological association with other anogenital and head and neck cancers (Garbuglia 2014). HPV16 (a member of *Alphapapillomavirus-9* species) is one of the most common HR types (Crow 2012).

Intracellular oncoproteins (E6 and E7) play an important role in the immortalization and malignant transformation of HPV-infected cells (Ghittoni, 2010). Eventhough both proteins are important for carcinogenesis, studying them independently allows for identification of pathways targeted by each oncoprotein. For the past 30 years, researchers have extensively studied how the E6 protein deregulates and transforms host cells (Klingelhutz, Roman 2012, Vande Pol, Klingelhutz 2013). HPV16 E6 confers viral persistence by down-regulating type I interferons (IFNs) alpha, beta (Ronco et al. 1998, Nees et al. 2001), kappa (Rincon-Orozco et al. 2009, DeCarlo et al. 2010, Reiser et al. 2011) as well as type II IFN gamma (de Gruijl et al. 1999). E6 prevents retention of Langerhans cells in infected epithelia through the down-regulation of E-cadherin (Matthews et al. 2003). E6 induces cellular immortalization through simultaneous upregulation of hTERT, the catalytic subunit of the enzyme telomerase (Klingelhutz et al. 1996, Veldman et al. 2003, Gewin et al. 2004, Katzenellenbogen et al. 2007), and degradation of the tumour suppressor protein P53 (Scheffner et al. 1993, Zanier et al. 2012). A transformed cellular phenotype is promoted when the C-terminus of E6 binds to PDZ domain-containing proteins, such as membrane-associated guanylate kinase (MAGI)-1 (Kranjec, Banks 2011), MAGI-2, MAGI-3 (Thomas et al. 2002), hScribble (Nakagawa, Huibregtse 2000) and hDlg (Kiyono et al. 1997, Gardiol et al. 1999), disrupting cell-cell signalling, cellular adhesion and cell polarity.

HPV16 is comprised of four sub-lineages or “variants”: A (European prototype sub-lineages A1/A2/A3 and the Asian sub-lineage A4), B (African-1 sub-lineages B1/B2), C (African-2) and D (North American sub-lineage D1 and Asian-American sub-lineages D2/D3) (Chen et al. 2005, Burk et al. 2013). We and other groups have been studying three HPV16 E6 protein variants (Figure 1): European prototype (EP), European-T350G (E-T350G: has an L83V single nucleotide polymorphism (SNP) in the E6 protein) and Asian-American (AA: has Q14H/H78Y/L83V SNPs in the E6 protein) (Zehbe et al. 1998, 2009, Zacapala-Gómez et al. 2016, Hochmann et al. 2019). Epidemiologically, the AA variant is associated with a higher risk for developing high grade cervical lesions and cervical cancer, as well as with an earlier cancer onset (Xi et al. 1997, Villa et al. 2000, Berumen et al. 2001, Xi et al. 2007). The epidemiological evidence surrounding the E-T350G variant is more complex, showing population-specific risk (Togtema et al. 2015, Zhang et al. 2016). Our functional investigations demonstrated that EP, E-T350G, or AAE6 retrovirally transduced into primary human foreskin keratinocytes (PHFKS) is able to immortalize these cells even in the absence of E7 (Niccoli et al. 2012, Togtema et al. 2015). While E-T350G and AAE6 promoted quicker cell growth and an earlier escape from growth crises than EPE6, only AAE6-transduced cells were able to form viable colonies in soft agar and demonstrate the ability to migrate and become invasive. Similar results were seen when E7 was also present (Richard et al. 2010). When keratinocytes were grown in 3D raft cultures, E-T350G and AAE6 caused higher grade dysplasia with more severe abrogation of differentiation patterns than EPE6 (Zehbe et al. 2009, Jackson et al. 2014). We found that AAE6 elevated the levels of glycolytic enzymes related to the tumour cell hallmark Warburg effect (Richard et al. 2010) and that of hypoxia inducible factor 1 alpha (HIF-1α) (Cuninghame et al. 2017), possibly contributing to AAE6’s enhanced proliferative abilities. The deregulation of HIF-1α was caused by AAE6’s ability to augment mitogen-activated protein kinase/extracellular related kinase (MAPK/ERK) signaling. It has also been shown that E-T350G differentially affects the amount of MAPK and phosphoinositide 3-kinases (PI3K)/activated kinase tyrosine (AKT) signaling (Chakrabarti et al. 2004, Sichero et al. 2012, Hochmann et al. 2016), binding to calcium-induced binding protein and hDlg (Lichtig et al. 2006) as well as E-cadherin (Togtema et al. 2015).

**Figure 1.**
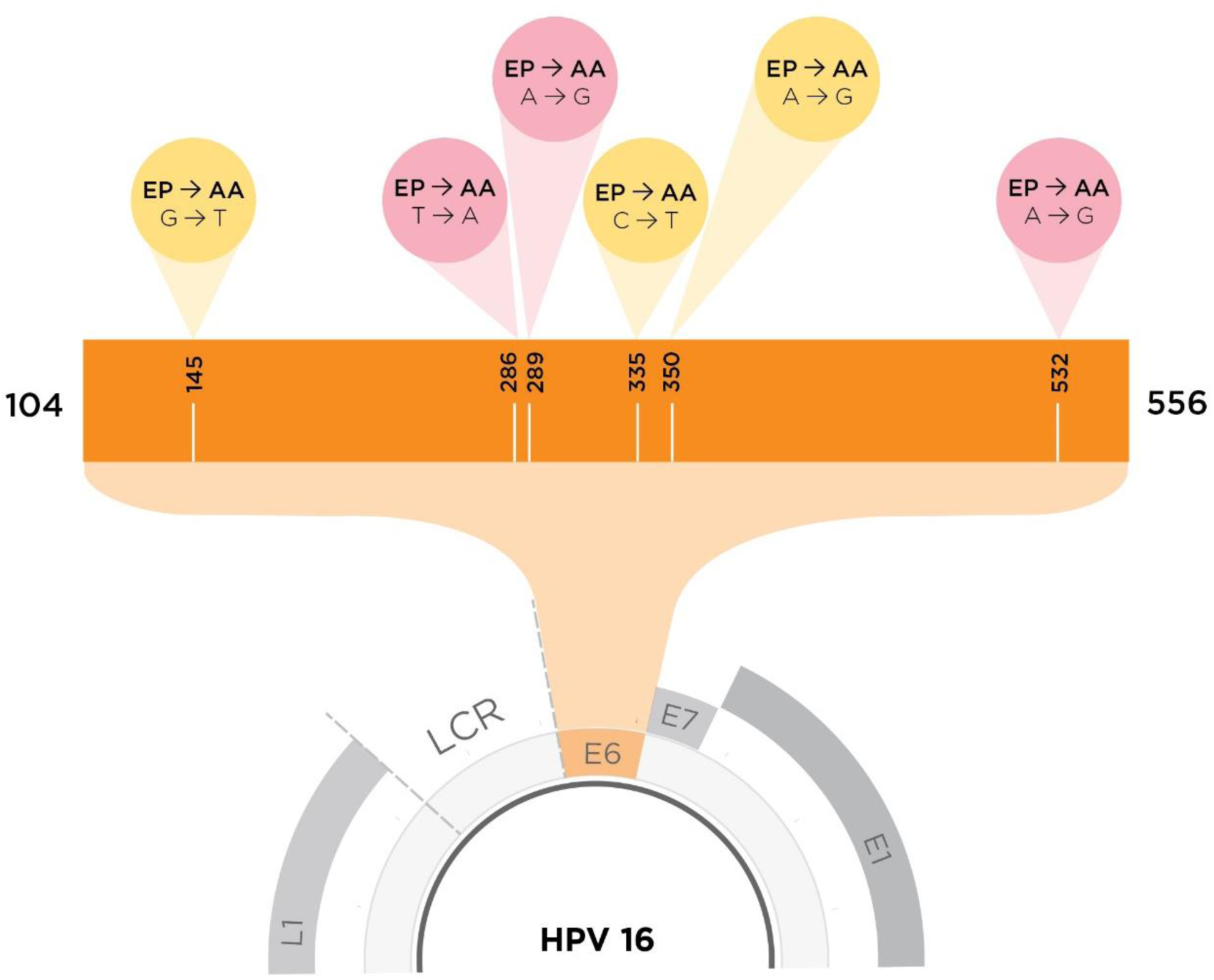
To scale depiction of HPV16 E6 SNPs between AAE6 and EPE6. The yellow cones indicate a SNP that results in an amino acid change (missense mutation), whereas the pink cones indicate no amino acid changes at that particular site (nonsense mutation). SNPs resulting in amino acid changes from EPE6 to AAE6 (Q14H, H78Y, and L83V) are found at nucleotide (NT) positions G145T, C335T, and T350G. SNPs that do not result in any change in amino acids are found at NT positions: T286A, A289G, and G532A (Zehbe et al. 1998).

To elucidate the mechanisms behind some of the above-mentioned functional observations, Zacapala-Gómez et al. (2016) completed a global transcriptome study and found 387 differentially-expressed genes in C33A cells and compared to C33A cells transfected with EPE6. The genes involved were related to cellular processes such as adhesion, angiogenesis, apoptosis, differentiation, cell cycle, proliferation, transcription and protein translation; specifically, they found over-representation of more than 1.5-fold for immunological processes for AAE6 compared to EPE6-transfected cells. Of course, the carcinogenic C33A cells may already have had underlying alterations in the expression of these pathways prior to the introduction of the E6 protein. A whole HPV16 genome sequencing study using 7116 pooled patient samples reported that the D2/3 (AA) as well as A3/4 sub-lineages show increased cervical cancer risk compared to A1 [EP of the current study] (Clifford et al. 2019). Thus, variant E6 proteins are not merely interacting in different amounts with the same set of host proteins. Instead, amino acid changes in E6 may allow it to bind a unique set of host proteins.

In this study, we explored the most likely mechanism for the described E6 functions: protein-protein interaction (PPI) between two key E6 variants and host cellular targets. We first surveyed the literature and summarized the findings of 50 previously reported HPV16 E6 interactors (Supplemental Table 1). Notably, most if not all, interaction experiments of the 50 known binders did not specify which HPV16 E6 sub-lineage was used other than “wildtype” or “prototype” with or without artificial mutants. Thus, we started filling this gap by investigating two common, naturally occurring E6 variants of the A1 (EPE6) and D2/D3 (AAE6) sub-lineages, based on our hypothesis that the type of PPIs between E6 and host cell proteins affects the carcinogenic potential of the HPV16 variant. Our methodological approach used primary human foreskin keratinocytes—naturally targeted by HPV—as well as state-of-the-art liquid chromatography-MS for virus-host PPI analysis and regularly-curated, freely-accessible databases. The obtained results will facilitate many future studies in HR HPV biology and may prove useful for anti-cancer treatment strategies targeting E6 (Togtema et al. 2019).

## Methodological Approach

### Wet lab methods

Cell culture, cell lysis, Western blot and co-immunoprecipitation (co-IP) methods used in this investigation are described in detail in Supplemental Data. The most challenging step in optimizing co-IP was protein elution, which was limited by several factors: compatibility in downstream liquid chromatography tandem mass spectrometry (LC-MS/MS) applications, the effectiveness of removing target proteins and minimizing antibody leeching. Elution methods described by other groups (White et al. 2012, Grace, Munger 2017) didn’t elute detectable levels of 16E6 in every replicate. Acidic elution was chosen for this study, since it was most effective at elution and also compatible with LC-MS/MS.

### Liquid chromatography tandem mass spectrometry (LC-MS/MS)

Two independent mass spectrometry trials were conducted with the two hemagglutinin-tagged (HA) variants: AAE6-HA, EPE6-HA and HA vehicle-transduced primary human foreskin keratinocytes (PHFK-HA) treated with the proteasome inhibitor MG132-DMSO or DMSO only, to yield the highest possible number of E6-binding partners (White et al. 2012). Following co-IP and elution, all samples were shipped to the Harvard Center for Mass Spectrometry (HCMS), Cambridge, MA, USA and processed as a paid service on demand. Detailed information can be found in Supplemental Data.

### Databases used for contaminant removal and identification of cellular targets

To determine the relationships between proteins identified by LC-MS/MS, we used three regularly-curated and freely-accessible databases: the Universal Protein Resource (UniProt) (The UniProt Consortium 2019) for molecular functions and biological processes; the Biological General Repository for Interaction Datasets (BioGRID) (Oughtred et al. 2019) for PPI analysis; and the Reactome Pathway Database (Reactome) (Jassal et al. 2019) to map E6 protein binders to the biological pathway in which they are involved. UniProt (whose protein abbreviations we adopted and reported in our study) provides an accession number (shown in our results) for each protein, which can be inserted into Reactome or BioGRID. Reactome generates a ranking of the 25 most significant pathways, meaning that these pathways are over-represented compared to others based on their probability value (P-value) and false discovery rate (FDR) (Fabregat et al. 2017). In our context, Reactome was used to discriminate non-specific binders and to determine how cellular pathways are [differentially] targeted by AAE6 vs. EPE6 and BioGRID if proteins of interest were interacting with each other.

### Post LC-MS/MS data contaminant removal

We merged the original “raw” HCMS Excel files that listed each trial (T1 and T2), condition (DMSO and MG132), E6 variant (AAE6 or EPE6) and PHFK-HA. After manual removal of sample-processing contaminants labelled *CON* (n=50)] from the merged Excel spreadsheet, we obtained a raw heatmap with a total of 1,584 proteins listed in alphabetical order (Supplemental Table 2). We then cleaned the data of ribosomal proteins (n=95) by running them through a Reactome (Fabregat et al. 2017) analysis using all the remaining proteins, then further removed proteins involved in pathways of RNA metabolism (n=89), yielding 1,464 remaining proteins (Supplemental Table 3). Proteins present in any of the PHFK-HA samples were likely to be contaminants and were likewise removed from the Excel spreadsheet (n=586; Supplemental Table 3).

### E6 interactors selection

#### Peptide method

In HPV-related publications where multiple Co-IP coupled to MS experiments were performed, two commomly used strategies for protein selection were reported: 2-peptide selection strategy or selection of proteins identified in more than one trial (Katzenellenbogen et al. 2007, White et al. 2012). The peptide method we used to select potential candidates among the remaining proteins followed one of these two stringent rules: any EPE6- or AAE6-targeted protein must have a sum of at least 2 peptides in either trial, independent of treatment, OR any EPE6- or AAE6-targeted protein must have at least 1 peptide in both trials, independent of treatment. After sorting the proteins with Excel, our peptide selection strategy left us with 25 cellular protein targets: 13 targeted by both variants and 6 unique to each AAE6 and EPE6 (Supplemental Table 4).

#### Protein-pathway method

We also designed a more inclusive approach to select additional potential E6 targets. We reasoned that, although it is possible that LC-MS/MS will miss several proteins in a given sample, it is far less likely that a whole pathway will be missed. Emphasis was on differences in proteins between 2 groups rather than on peptide abundance alone, i.e. AAE6 versus PHFK-HA, EPE6 versus PHFK-HA and AAE6 versus EPE6. We detected 498 proteins in AAE6, 380 in EPE6 (n=108 overlap) and 586 proteins in PHFK-HA samples (Supplemental Table 3). Each group of proteins was then analysed using Reactome to identify the pathway involving these proteins: 251/498 AAE6-targeted proteins were matched to 861 pathways, 199/380 EPE6-targeted proteins to 794 pathways and 323/586 “unspecific” protein binders in PHFK-HA were matched to 954 pathways.

We then compared the pathways obtained for each group and selected those that aligned with AAE6 and/or EPE6 interactors but not with PHFK-HA interactors. We reasoned that pathways only targeted by AAE6 or EPE6 would be most specific, leaving us with 90 unique proteins for AAE6, 61 unique proteins for EPE6 and 20 additional proteins common to both, i.e. altogether 171 potential E6 binders (Supplemental Table 5). These proteins were then run independently (all AAE6 interactors and all EPE6 interactors separately) on Reactome to assess which pathways were significantly over-represented for each E6 sub-lineage.

## Results and Discussion

### PEPTIDE METHOD: E6 activities relate to P53 tumour suppression, hypoxia, energetics, chromosome remodeling and innate immunity

Our *peptide method* generated 25 candidates (6 unique AAE6 and 6 unique EPE6 interacting proteins along with 13 common E6-interacting proteins between AAE6 and EPE6; Figure 2A, Supplemental Table 4). These potential E6 interactors were screened for functions related to HPV16-related tumourigenesis and immune suppression, resulting in a short-list of 7 proteins. Three proteins are targeted by AAE6: inositol (INO)80 complex subunit B (IN80B, Q9C086), ribosomal protein S6 kinase alpha-4 (KS6A4, O75676), and prokineticin-2 (PROK2, Q9HC23). One is targeted by EPE6: interferon-induced GTP-binding protein (MX2, P20592). Three are targeted by both AAE6 and EPE6: E3 ubiquitin-protein ligase trip12 (TRIPC, Q14669), nucleolar GTP-binding protein 2 (NOG2, Q13823), charged multi-vesicular body protein 4b (CHM4B, Q9H444).

**Figure 2.**
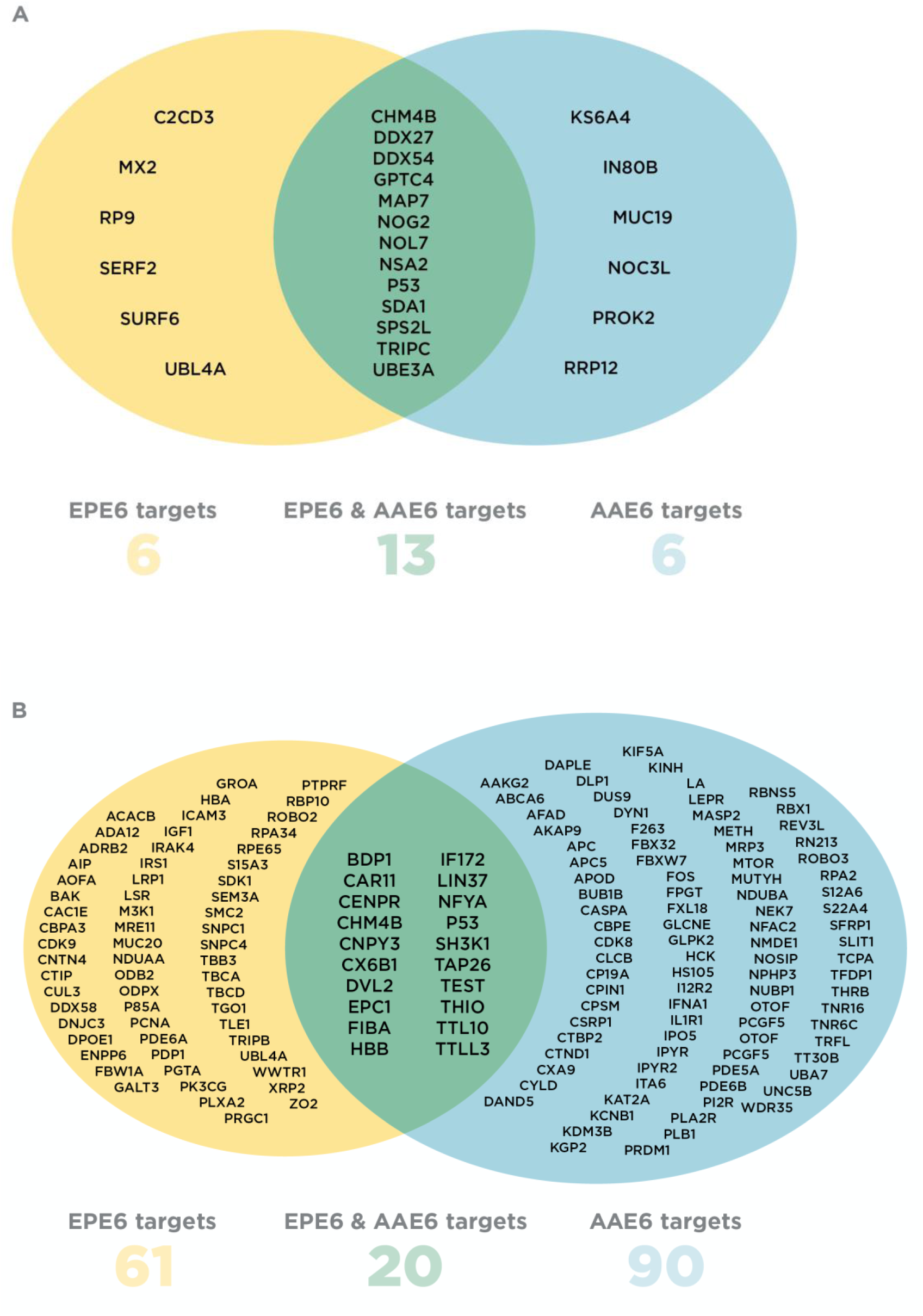
Venn diagrams representing AAE6 and EPE6 host cellular targets for both selection approaches: Peptide method (A) and Protein-pathway method (B). EPE6-specific interactors are in yellow, AAE6-specific interactors are in blue and proteins common to both variants are in green. Protein names are abbreviated based on UniProt protein rather than gene nomenclature, enabling easy identification for searches and further information.

#### AAE6 affects chromosome remodeling, hypoxia and innate immunity

**IN80B**, an ATPase, is part of the ATP-dependent INO80 remodeling complex containing at least 12 other proteins. The INO80 complex functions in transcriptional regulation, DNA replication and repair, telomere maintenance and chromosome segregation (Min et al. 2013). In a mouse model of cervical cancer (Hu et al. 2016), INO80 was overexpressed and, when bound to the Nanog transcription start site, this transcription factor’s expression was enhanced. Blocking its interaction led to a decrease in proliferation. In addition, yeast 2 hybrid (Y2H) experiments indicated that IN80B interacts with HPV16 E7 and HPV5 E7 (Rozenblatt-Rosen et al. 2012), suggesting that both E6 and E7 could alter the function of IN80B. Proximity label MS showed that the Myc proto-oncogene protein (MYC), a known target of E6 (Gross-Mesilaty et al. 1998, Supplemental Table 1), also interacts with NOC3L and RRP12 (Kalkat et al. 2018, Supplemental Table 1). Notably, IN80B and RRP12 were present only in the AAE6 sample, NOC3L was targeted by both E6 variant proteins, whereas RP9 was unique to EPE6. Hence, even if EPE6 does not immunoprecipitate IN80B, it may alter its functions indirectly. Reactome analysis showed IN80B to be associated mainly with DNA damage and repair pathways and metabolism of proteins (Supplemental Table 6).

**KS6A4** was initially identified as mitogen- and stress-activated protein kinase 2, **MSK2** (75% identical to MSK1) and is activated by the mitogen activated protein kinase (MAPK)/extracellular signal-regulated kinase (ERK) or stress-activated protein kinase (SAPK)/p38 phosphorylation cascade (Deak et al. 1998; reviewed in Wiggin et al. 2002). MSK1 and 2 regulate the transcription factors cAMP response element-binding protein (CREB) and activating transcription factor 1 (ATF1) through phosphorylation (Deak et al. 1998). MSKs were suggested to be negative regulators of the innate immune system as their regulation of CREB and ATF1 also controls the expression of interleukin-10 and dual specificity protein phosphatase 1 (Ananieva et al. 2008). They are further involved in chromatin remodeling by phosphorylating histone H3 and the high mobility group chromosomal protein (HMG) 14 (Soloaga et al. 2003). Chromatin remodelers are “essential for all DNA-dependent processes” (Längst, Manelyte 2015)—a fact that connects them with the above-described AAE6 interactor IN80B. MSK2 negatively regulates p53 in a kinase-independent manner through its interaction with P300 and its binding to the NOXA (DNA damage- and genotoxic stress-related) promoter (Llanos et al. 2009). Consistent with these inhibitory functions, MSK2 inhibits apoptotic processes (Llanos et al. 2009). It is overexpressed in squamous cell carcinomas of the cervix. Its knockdown was reported to inhibit the phosphorylation of paired-box gene 8 (PAX8) and the retinoblastoma protein (pRB), as well as to suppress cell cycle-stimulating factors E2F1 and cyclin A2, leading to a decrease in SiHa and HeLa cell proliferation (Wu et al. 2019). Stabilizing MSK2 by AAE6 could complement the well-known action of E7 in causing the release of E2F1 through binding to pRB (Sherr, Roberts 1999).

We have previously identified increased signaling in the ERK1/2 pathway for AAE6 compared to EPE6, suggesting that MSK2 is a key contributing factor to increased hypoxia-inducible factor (HIF)-1α seen in AAE6 over EPE6 cells in the hypoxic tumour environment (Cuninghame et al. 2017). Interestingly, MSK2 has been shown to activate NF-κB/p65 under different stimuli (Vermeulen et al. 2009). It is also known that under hypoxic conditions, NF-κB/p65 is a regulator of HIF-1α gene expression (Yoshida et al. 2013). Although it has not been shown that MSK2 activates NF-κB/p65 under hypoxic conditions, one could speculate that ERK1/2 activation of MSK2 promotes NF-κB/p65 positive regulation of HIF-1α. Consistent with this notion, MSK2 was only targeted by AAE6. Others have likewise observed that ERK1/2 phosphorylation of MSK2 is higher in AAE6 compared to EPE6 and E-T350G (also called L83V) (Hochmann et al. 2016). We conclude that the AAE6 PPI with this kinase may be one of the underlying factors causing the differential finding in sugar metabolism between AAE6- and EPE6-transduced PHFKs previously reported by us (Richard et al. 2010; Cuninghame et al. 2017). Notably, Reactome did not yield pathways related to hypoxia with MSK2 (Supplemental Table 6).

**PROK2** along with PROK1 are chemokine-like proteins (attracting leukocytes to an inflammatory site) usually expressed by components of the innate immune system, such as macrophages with specific roles in host defence and angiogenesis during virus-related cancers (Lauttia et al. 2014). They are ligands transducing their signals through G protein–coupled receptors (Zhou et al. 2006). PROK2 expression is upregulated in cases of human Merkel cell carcinoma (MCC) containing the MC polyomavirus (MCpyV) (Lauttia et al. 2014). MCpyV shares similarity with HPV via the large T antigen and E6, respectively, targeting P53. This, and the augmentation of tumour-infiltrating macrophages, resulted in positive survival outcomes while the opposite result was obtained with increased PROK1 expression and absence of MCpyV. Using a colorectal cancer cell and mouse model, PROK2 promoted angiogenesis, leading to an increase in colon tumour mass (Kurebayashi et al. 2015). Interestingly, PROK2 is known to sequester the promoter of HIF-1, alluding to the hypoxic tumour environment (resonating with MSK2 activities above) and to alter extracellular matrix potentially controlling angiogenesis (LeCouter et al. 2001, 2003). PROK2 was found in 7 Reactome pathways mostly related to receptor-mediated signal transduction but not to hypoxia—each associated with AAE6 and EPE6, albeit with different entities (Supplemental Table 6). Taken together, PROK2 may be another interesting AAE6-target, as its dual role in HIF-1 signaling and immune biology may contribute to the “success” of this sub-lineage in tumour development.

#### EPE6 interferes with host-cellular immune surveillance

Mx GTPases MX1 and **MX2** are “dynamin-like antiviral machines of innate immunity” (Haller et al. 2015). They are essential components of the antiviral response induced by type I and III interferon (IFN) and act as inhibitors of early viral replication (Haller et al. 2015). Two MX2 isoforms (with or without nuclear localisation signal) are found in the nucleus or cytoplasm where they interact with different key viral components. For example, during HIV infection, MX2 targets viral genome uncoating, nuclear uptake and integrase activity of the pre-integration complex. MX2 also has a potential function in cervical carcinogenesis. In the HPV16-positive cell line W12, the selection of infected keratinocytes with integrated viral DNA requires the loss of the episomal genome and that episomal expression of E2 could limit transcription of the integrated viral DNA (Pett et al. 2006). Microarray analysis revealed that episome loss was associated with the expression of type I IFN pathway-inducible genes including MX1 and 2 (Pett et al. 2006). Interaction of EPE6 with MX2 and the subsequent potential alteration of MX2 activity could therefore be a factor explaining the different genome integration capacity of AAE6. Based on Y2H experiments, MX2 interacts with the histone-lysine N-methyltransferase EHMT2 (Rolland et al. 2014), which increases P53-dependent expression of pro-apoptotic genes, i.e. Bax and PUMA (Rada et al. 2017). EHMT2 is also a known interactor of EP300/CBP (Rada et al. 2017), which in turn is a known E6 binder (above and Supplemental Table 1). Reactome pathways for both AAE6 and EPE6 were related to IFN signaling and antiviral mechanisms but with different interactors (Supplemental Table 6).

#### AAE6 and EPE6 both disrupt P53 activities

**NOG2**—a GTPase—is involved in the regulation of G1 to S phase transition (Datta et al. 2015) and ribosomal biogenesis (Essers et al. 2014). NOG2 induces cell proliferation by increasing TP53 (and its downstream product, CDKN1A/p21) protein levels, and decreasing RPL23A protein levels (Datta et al. 2015). Interest in NOG2 arose from the observation that the protein is overexpressed in some cancers, such as breast or colorectal (Racevskis et al. 1996). NOG2 induces P53 activation by inhibiting RPL23A (60S ribosomal protein L23a), a ribosomal protein that triggers p53 degradation via Mdm2. P53 expression leads to the expression of the cyclin-dependent kinase inhibitor p21 (Datta et al. 2015), which is required for the formation of the cyclin D1-CDK4 complex. At higher concentrations, p21 inhibits the cyclin D1-CDK4 complex (LaBaer et al. 1997). An activated cyclin D1-CDK4 complex leads to a decrease in pRB phosphorylation that, in turn, releases E2F1 to promote cell proliferation (Datta et al. 2015). NOG2 is a known interactor of IN80B (Cloutier et al. 2017, Supplemental Data) and both proteins interact with RPL23A (Datta et al. 2015, Cloutier et al. 2017). In addition, NOG2 interacts with MYC (Kalkat et al. 2018) which is involved in P53 stabilization. Taken together, these data suggest that NOG2 is involved in several pathways that could all affect P53 activity in the HPV context. An interactome study indicated that in addition to IN80B, NOG2 also interacts with two other proteins selected by the peptide method, i.e. ribosome biogenesis protein NSA2 homolog (Nop seven-associated 2, NSA2) (Huttlin et al. 2015) and G patch domain-containing protein 4 (GPTC4) (Huttlin et al. 2017) (Supplemental Table 4). These data originate from large interactome studies and the consequences of each interaction were not investigated (Huttlin et al. 2015, 2017). GPTC4 functions are largely unknown, nevertheless, information available on NSA2 indicates that the protein could be involved in the same pathways as NOG2 as they appeared to have similar functions. No Reactome pathways could be aligned with NOG2 (Supplemental Table 6).

**TRIPC** is an E3 ubiquitin-protein ligase that shares similarities with UBE3A (E6AP). It contains the conserved homologues to E6AP carboxy terminus (HECT) domain and multiple LxxLL (where x denotes any amino acid) motifs that correspond to the E6 binding site on E6AP (Vande Pol, Klingelhutz 2013, Zanier et al. 2013, Larrieu et al. 2020). There are 4 motifs starting at various residues: LQALL (402), LITLL (485), LHFLL (697) and LDQLL (1862) present throughout TRIPC that could potentially allow interaction with E6. TRIPC triggers the ubiquitination and degradation of several proteins including the P53 activator protein ADP-ribosylation factor (ARF, p14 in humans) (Haupt et al. 1997, Collado, Serrano 2010) potentially doubling the effect of P53 inactivation by E6 as ARF acts upstream of E6AP-mediated P53 degradation. Interestingly, TRIPC-dependant ARF degradation is inactivated by MYC or tumour necrosis factor receptor type 1-associated DEATH domain protein (TRADD) binding to TRIPC (Chen et al. 2010, Chio et al. 2012). Since MYC is a known E6 target, E6 binding to TRIPC could promote ARF degradation by stimulating TRIPC activity and E6 binding to TRIPC and/or MYC could protect TRIPC from MYC inactivation. In addition to ARF, the Brahma-related gene 1-associated factor 57 (BAF57) is another protein targeted by TRIPC. BAF57 degradation by TRIPC could be inhibited when BAF155 is bound to TRIPC (Keppler, Archer 2010). BAF57 is a canonical component alongside BAF53 of the SWItch/sucrose non-fermentable (SWI/SNF) chromatin remodeling complex (Chen, Archer 2005)—first detected in yeast (Cristofaro et al. 2001). Interestingly, BAF53 is essential for the expression of E6 and E7 when the viral genome has been integrated into the host cell (Lee et al. 2011). TRIPC is involved in Reactome pathways mainly associated with Antigen processing, Ubiquitination and Immune system (Supplemental Table 6). Interestingly, E6AP is found in all the pathways involving TRIPC and all pathways involving TRIPC are connected to AAE6 and EPE6 but with different entities (Supplemental Table 6).

**CHM4B** is thought to be a subunit of the endosomal sorting complex required for transport (ESCRT)-III complex involved in the formation of multi-vesicular bodies and is important in membrane fission processes, including the budding of enveloped viruses (Strack et al. 2003, Hu et al. 2015). CHM4B is overexpressed in hepatocellular carcinomas (Hu et al. 2015) and some head and neck squamous cell carcinomas (Gollin 2014). Co-IP experiments indicated that CHM4B interacts with the inhibitory P53 isoform Δ133p53α that can block the activity of full-length P53 (Horikawa et al. 2014). Other interesting interactors of CHM4B are interferon-responsive factor (IRF)-2 (Hubel et al. 2019), breast cancer-associated protein (BRCA) 2 (Malik et al. 2016) and E-cadherin (Guo et al. 2014). CHM4B presence in Reactome is mostly associated with Antigen processing, HIF viral life cycle and Autophagy, some of which are shared among both variants while others differ (Supplemental Table 6): e.g. Late endosomal microautophagy pathways also include Hemoglobin subunit beta HBB (P68871) [AAE6 and EPE6], as well as HIV infection pathways also include E3 ubiquitin-protein ligase RBX1 (P62877), Tyrosine-protein kinase HCK (P08631) [AAE6], Cyclin-dependent kinase 9 CDK9 (P50750) and F-box/WD repeat-containing protein 1A FBW1A (Q9Y297) [EPE6].

### PROTEIN-PATHWAY METHOD: AAE6 is more “successful” than EPE6 in the malignant transformation process

AAE6 (n=90), EPE6 (n=61) and overlapping AAE6/EPE6 binders (n=20) selected by the *protein-pathway method* (Figure 2B; Supplemental Table 5) were each collectively analyzed using Reactome. Within the 25 most significant pathways, 26/90 entities were identified for AAE6 (Supplemental Table 7) and 48/61 for EPE6 (Supplemental Table 8). Out of 20 overlapping AAE6/EPE6 targets, 5 were identified within the 25 most significant pathways targeted by both variants, whereas 8 were only seen in EPE6-targeted pathways (Supplemental Table 9). While the P-value was significant throughout for both sub-lineages and the overlapping proteins, the FDR was statistically significant (P<0.05) in 18/25 pathways for AAE6 only (Supplemental Tables 7, 8 and 9). Nevertheless, for comparison, we investigated results of both E6 sub-lineages and overlapping E6 targets. AAE6 and EPE6 entities belonging to these pathways are also listed.

#### AAE6 affects Notch signaling, hypoxia, metabolism and DNA base excision repair

AAE6 was strongly associated with 11 Notch1 signaling pathways (**1**,**2**,**3**,**9**,**10**,**11**,**12**,**13**,**15**,**16**,22) with the same protein targets (Supplemental Table 7): cyclin-dependent kinase 8 (CDK8, P49336), histone acetyltransferase KAT2A (Q92830), E3 ubiquitin-protein ligase RBX1 (P62877) and two isoforms of the F-box/WD repeat-containing protein 7 (FBXW7, Q969H0-1, Q969H0-4). CDK8 phosphorylates the neurogenic locus notch homolog protein 1 (NOTC1), targeting it for ubiquitination and degradation through the proteasome. Consequently, NOTC1 is not acetylated by mastermind-like protein 1 (MAML1) and its transcription is not enhanced by EP300 (Popko-Scibor et al. 2011). Notch signaling regulates cell homeostasis (balancing proliferation, differentiation and survival/apoptosis), while abberant Notch signaling seems to be a major factor in driving epithelial neoplasia. Indeed, it was recently reported that “Rare driver mutations in head and neck squamous cell carcinomas converge on Notch signaling” (Loganathan et al. 2020). RBX1 and FBXW7 are also part of Wnt signaling (pathway **14**), suggesting communication between these two pathways. Wnt/beta catenin signaling plays a role in development and adult homeostasis. In the latter, it is mostly inactive and controlled by kinases, such as glycogen synthase kinase 3 (GSK3), casein kinase 1 (CK1), axin and adenomatous polyposis coli (APC), a tumour suppressor gene often mutated in colon cancer (Verheyen, Gottardi 2009 and references therein). Interestingly, using our peptide method, APC (P25054) was found only once with just one unique peptide, yet it appeared in pathway **14** targeted by AAE6.

We also found AAE6 binders belonging to defective base excision repair (BER) associated with the human homologue of the Escherichia coli mutY gene (hMYH), i.e. two isoforms of its encoded protein adenine DNA glycosylase (MUTYH): Q9UIF7-3 and −6 (pathways **5**,**7**,19). MUTYH germline mutations of BER (pathways **5**,**7**,19) cause MUTYH-associated polyposis (MAP), a disorder similar to familial adenomatous polyposis (FAP), caused by mutations in the APC gene (Mazzei et al. 2013). This association between AAE6 and BER indicates that the AA (D2/D3) sub-lineage integrates earlier into the host genome than EP (A1), supported by observations in an organotypic cell culture model of early cervical carcinogenesis (Jackson et al. 2014, 2016). Indeed, another group reported that BER is essential for the HIV provirus DNA to integrate into the host genome, proposing an interesting analogy with transposable elements (Yoder et al. 2011). In “The landscape of viral association with human cancers” study led by the Pancancer Analysis of Whole Genomes Consortium (Zapatka et al. 2020), HPV16 integration seems to be associated with fragile sites or regions with limited access to DNA repair complexes. Our findings from the peptide and protein method corroborate these results: ATP-dependent chromosome remodeling complex INO80 implicated in DNA repair enhances BER activity (Hinz, Czaja 2015), and the antiviral function of MX2 resulting in viral integration (Pett et al. 2006) would “help” AAE6 but not EPE6 to insert itself into the host cell genome.

We observed AAE6 binding to the following P53-regulated metabolic genes (pathway **8**): P53 (P04637), MTOR (P42345), thioredoxin (THIO, P10599), trinucleotide repeat-containing gene 6C protein (TNR6C, Q9HCJ0), cytochrome c oxidase subunit 6B1 (CX6B1, P14854) and 5’-AMP-activated protein kinase subunit gamma-2 (AAKG2, Q9UGJ0). Note that P53-regulated metabolic genes also communicate with Wnt signaling via AAKG2. Hence, AAE6-targeted cellular proteins derive from the axis of Wnt and Notch1 signaling, as well as Wnt signaling and P53-regulated metabolic genes. The most striking candidates for this axis are RBX1, FBXW7, KAT2A and CDK8, due to their presence in 8 to 11 pathways (Supplemental Table 7).

AAE6 targeting of pathway **8** proteins (and its association with MSK2, as discovered in the *peptide method*) explains how this variant deregulates cellular metabolism via the Warburg effect (Richard et al. 2010, Cuninghame et al. 2017) and promotes a hypoxic environment via elevated HIF-1α levels (Cuninghame et al. 2017). Deregulated energetics and hypoxia—two hallmarks of cancer (Hanahan, Weinberg 2011)—may cause the “higher” carcinogenic ability of AAE6. While EPE6 also targets P53, CX6B1 and THIO of pathway **8**, AAE6 may have an advantage over EPE6 due to the additional actions of MTOR, TNR6C and AAKG2. Moreover, RBX1, FBXW7 and CDK8 (Notch1 signaling pathways **1**,**2**,**3**,**9**,**10**,**11**,**12**,**13**,**15**,**16**), as well as MTOR (pathway **8**), are known HIF-1α binders (Galbraith et al. 2013, Heir et al. 2013, Perez-Perri et al. 2016), reinforcing AAE6’s effect on hypoxia. Indeed, alteration of MTOR function is known to promote HIF-1α translation (Knaup et al. 2009). CDK8 and its associated protein mediator of RNA polymerase II transcription subunit 1 and 26 are recruited by the HIF-1α transactivation domain at the promoter of several hypoxia-inducible genes, leading to the activation of the super elongation complex (SEC) to promote RNA polymerase II (RNAPII) elongation (Galbraith et al. 2013). In addition, proto-oncogenes c-Fos-encoded protein (FOS, P01100) (pathways 21,24) and 6-phosphofructo-2-kinase/fructose-2,6-bisphosphatase 3 (F263, Q16875), were found to be associated with HIF-1α, despite not directly interacting with it (Onnis et al. 2013, Kumar et al. 2015). HIF-1α and FOS both bind the vascular endothelial growth factor promoter under hypoxic conditions (Onnis et al. 2013). F263, a master regulator of glycolysis, is activated by HIF-1α and could be of importance for Warburg effect development (Minchenko et al. 2002).

Finally, Forkhead box protein O1 (FOXO)-mediated transcription (pathway 25 with AAE6 interactors and pathway 5 with proteins common to AAE6 and EPE6) overlaps the AAE6 and EPE6 variants via nuclear transcription factor Y subunit alpha NFYA (P23511) and THIO (Supplemental Table 9). AAE6 also interacts with the ATP-binding cassette sub-family A member 6 (ABCA6, Q8N139) and F-box only protein 32 (FBX32, Q969P5) which are involved in the FOXO-mediated transcription pathway, indicating that AAE6 could disrupt the pathway more efficiently than EPE6. Of interest, FOXO1 expression is decreased in CaSki and SiHa cells but the regulation of FOXO1 in cervical cancer is not yet fully understood (Zhang et al. 2015). FOXO transcription factors act in pathways controlling cell survival, growth, differentiation and metabolism in various scenarios, such as growth factor deprivation, starvation and oxidative stress (Eijkelenboom, Burgering 2013).

#### EPE6 is associated with DNA damage repair and cancer pathways

Up to 8 EPE6 interactors are part of homology directed repair (HDR) DNA damage pathways 5,9,19,22: P53, THIO, proliferating cell nuclear antigen (PCNA, P12004), CX6B1, double-strand break repair protein MRE11 (P49959), cyclin-dependent kinase 9 (CDK9, P50750), geranylgeranyl transferase type-2 subunit alpha (PGTA, Q92696) and DNA endonuclease RBBP8 (Q99708). PCNA also interacts with the adenine DNA glycosylase MUTYH (Parker et al. 2001), linking the two E6 variants. However, while some of the targeted pathways overlap, most of the entities associated with the individual E6 variant differ (Supplemental Table 6). Like AAE6, EPE6 seems to target various pathways crucial for carcinogenesis, such as Hippo, PI3K/AKT, growth factor initiated mesenchymal-epithelial signal transduction (MET) and epithelial growth factor receptor (EGFR) (pathways 1,3,7,10,14,24). Hippo communicates with the Wnt pathway through segment polarity protein dishevelled homologue DVL-2 (DVL2, O14641) via EPE6’s pathway 1 and AAE6’s pathways **14** and **17**. The most notable EPE6 target seems to be PI3K regulatory subunit alpha (P85A, P27986), detected in 5 pathways related to PI3K, MET and EGFR (3,7,10,14,24), which all communicate with each other (Supplemental Table 8). Interestingly, some HIF-1α binders were also found to be targeted with EPE6, i.e. WW domain-containing transcription regulator protein 1 (WWTR1, Q9GZV5) (pathway 1) and CDK9 (pathway 22) (Galbraith et al. 2013, Xiang et al. 2015). In breast cancer, WWTR1 (also called transcriptional co-activator with PDZ-binding motif) expression is activated by HIF-1α and is important for the maintenance of breast cancer stem cells (Xiang et al. 2014). Since CDK9 is a component of SEC recruited by CDK8 upon interaction with HIF-1α (Galbraith et al. 2013), both AAE6 and EPE6 could potentially alter RNAPII elongation. Nevertheless, only CDK8 is essential to the process (Galbraith et al. 2013) suggesting that AAE6 and EPE6 work differently to modify RNAPII elongation.

#### AAE6 and EPE6 both prevent P53 activity

The 25 most significant pathways generated for the 20 overlapping interactors of AAE6 and EPE6 are listed in Supplemental Table 9. No differential pathway enrichment was noted, as all 124 pathways showed the same FDR value of 0.09. Out of 20 overlapping proteins, 15 were distributed among the top 25 pathway, s with 13 found in the pathways described for AAE6 and EPE6: P53, NFYA, DVL2, THIO, CX6B1, CHM4B, HBB, inactive polyglycylase tubulin tyrosine ligase-like protein 10 (TTL10, Q6ZVT0), tubulin monoglycylase TTLL3 (Q9Y4R7), SH3 domain-containing kinase-binding protein 1 (SH3K1, Q96B97), fibrinogen alpha chain (FIBA, P02671), coiled-coil domain-containing protein 59 (TAP26, Q9P031) and testisin PRSS21 (Q9Y6M0). Consequently, most pathways identified with these proteins were involved in the same pathways as those described for AAE6 or EPE6 above, albeit with fewer entities (Supplemental Table 9). These results suggest that several pathways are targeted by both AAE6 and EPE6, but with different efficiencies (Supplemental Table 9). The two remaining proteins, protein lin-37 homologue (Q96GY3) and centromere protein R (Q13352), were found in pathway 14 (Cell cycle, Mitotic) alongside P53 and CHM4B. Overall, P53 is the main contributor found in 13 of the 25 pathways (1,4,7,10,12,13,14,18,19,20,23,24; Table 4), and is the only entity found in 9 of them. Thus, P53 appears to be the most important target between AAE6 and EPE6.

## CONCLUSIONS

This study is one of few reporting PPIs in a global perspective focusing on two common E6 sub-lineages with varying prevalence in cervical cancer. Using two different approaches to dissect the AAE6 and EPE6 variants and their cellular interators (Figure 3), it was clear that AAE6 takes the lead in targeting hypoxia and energetics, while both E6 proteins equally inactivate p53 by multiple means. The latter observation, along with the fact that AAE6 is also very active in subverting Notch1 signaling, are completely new discoveries and are strengthened by the finding that hypoxia requires Notch1 signaling to maintain the undifferentiated cell state in various stem and precursor cell populations (Gustafsson et al. 2005). The identification of the viral defence protein MX2 targeted by EPE6, making this variant less likely to integrate into the host cell genome than AAE6, bolsters our earlier *in vitro* data describing this phenomenon (Jackson et al. 2014, 2016). Targeting BER by AAE6 further underlines these findings. Subtle and indirect changes in viral immune evasion mechanisms were also uncovered for both variants. We are mindful that our results must be validated with additional wet lab experiments, such as reverse co-IP and RNA interference to elucidate gain or loss of function of reported candidate binders. Although only a small snapshot of identified proteins was provided here, confirming these new discoveries will provide vital information about E6’s ability to drive tumourigenic processes. Our study strengthens the notion that E6 is an excellent target for an anti-HPV treatment (Togtema et al. 2019), whose design would need to consider E6 sub-lineage differences.

**Figure 3.**
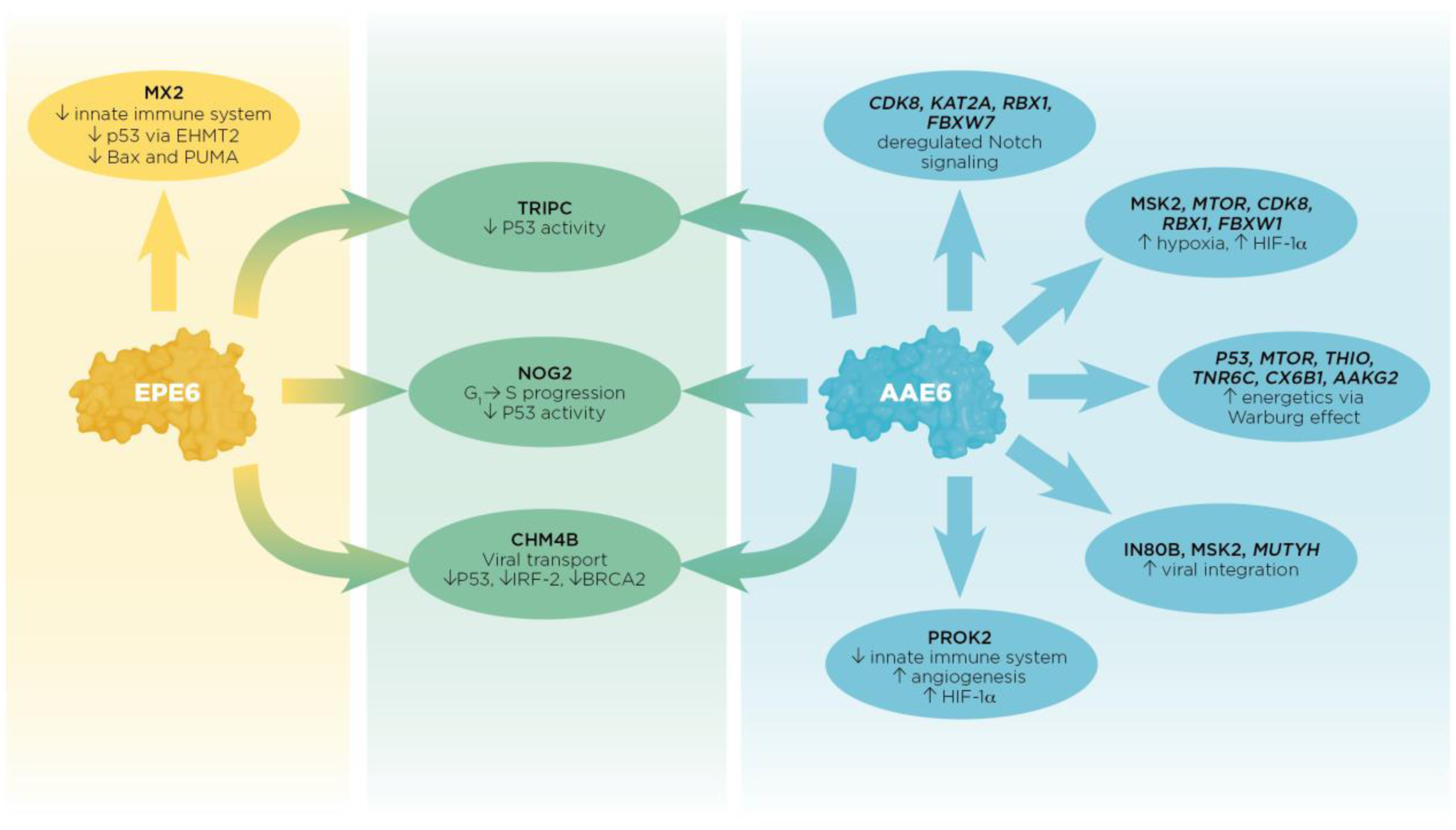
Summary of individual and tandem effects between AAE6 and EPE6 PPIs with host cellular proteins by which E6 may promote cell-transforming processes. Candidates from both approaches are shown for AAE6 only due to the lack of significant FDR values for EPE6 and AAE6/EPE6 combined using the Protein-pathway method (upright font=Peptide method, *italic* font= Protein-pathway method). Candidates from the Peptide method are shown for AAE6, EPE6 and AAE6/EPE6 (upright font). The first-described E6 binder (P53; Supplemental Table 1) is most likely targeted through several mechanisms shared by both E6 variants (middle). AAE6 and EPE6 proteins equally bound 3 proteins known to affect P53 inactivation: TRIPC, CHM4B, and NOG2. Collectively, these interactions could result in decreased P53 function far beyond its well-described E6- and E6AP-mediated degradation through the proteasome. Potential interactions unique to AAE6 (right) are mostly associated with Notch signaling, hypoxia, energetics, DNA base excision repair, and to some extent with the innate immune system. Notably, some AAE6-targeted molecules have multiple roles, e.g. MTOR is associated with hypoxia and metabolism, CDK8 with hypoxia and Notch signaling and MSK2 with hypoxia and chromatin remodeling further underlining this variant’s “lead” in the malignant process. AAE6’s association with MUTYH, IN80B and MSK2 could promote its integration potential and the binding to PROK2 could promote both angiogenesis and hypoxia within infected cells. Being a chemokine-like molecule, PROK2 is implicated in the innate immune system normally attracting macrophages to the site of inflammation. In the hypoxic and consequently more acidic tumour environment, tumour-associated macrophages (TAMs) may develop from original site-filtritating macrophages adapting to the tumour microenvironment. Tumour growth is then promoted by the positive feedback loop between TAMs and epithelial cells via the expression of colony-stimulating factor and epithelial growth factor, respectively. EPE6’s binding to MX2 (left) may limit this variant’s ability to integrate into the host genome effectively while it may be more successful in deregulating the host’s viral defence. MX2 also affects the P53 pathway, e.g. via EHMT2 and the expression of pro-apoptotic genes Bax and PUMA further duplicating well-established anti-P53 E6 activities.

## Supporting information

Dayer et al._Supplemental Data

Dayer et al._Supplemental Table 1

Dayer et al._Supplemental Table 2

Dayer et al._Supplemental Table 3

Dayer et al._Supplemental Table 4

Dayer et al._Supplemental Table 5

Dayer et al._Supplemental Table 6

Dayer et al._Supplemental Table 7-9

## Acknowledgements

This work was supported by a Natural Sciences and Engineering Research Council of Canada (NSERC) grant to I.Z. (NSERC Discovery Grant #RGPIN-2015-03855), a Mathematics of Information Technology and Complex Systems grant to G.D. (Mitacs Elevate Postdoctoral Fellowship IT15879) and an Ontario Graduate Scholarship to M.L.M. We are grateful to Megan Teghtmeyer for assisting with some wet lab experiments.

