## Supplementary material for "Virus-host protein-protein interactions between human papillomavirus 16 E6 A1 and D2/D3 sub-lineages: variances and similarities": Dayer et al._Supplemental Data

### Supplemental File: Wet Lab Methods

***Cell culture***

High-passage (at passage 60 or higher), primary human foreskin keratinocytes [PHFKs], previously transduced with either the AAE6 or EPE6 gene HA-tagged at the C-terminus (Niccoli et al., 2012) and control PHFKs transduced with the HA-tag alone were used. HA allows for highly specific IP of a target protein bound to an affinity tag (Free et al. 2009). It was placed at the C-terminus to reduce any disruption in interactions present at or near E6 SNP sites keeping in mind that this procedure could hinder the E6's potential to bind to PDZ-interacting proteins. Cells were cultured in complete keratinocyte growth medium (KGM, EpiLife medium completed with 60 µM calcium (EpiLife, Fisher Sci., Waltham MA, Cat# MEPI500CA); 1 X antibiotic/antimycotic solution (Fisher Sci., Cat# SV3007901) and 1 X human keratinocyte growth hormone (HKGS, Fisher Sci., Cat# S0015); 1.5 X 10^6 cells were seeded into Nunc™ EasYFlask™ 225 $cm^2$ culture flasks (Fisher Sci., Cat# 12-565-221) containing 33 mL KGM. Medium changes occurred every 48 hours and cells were grown until ~ 80% confluence. After trypsinization, cells were centrifuged at 750 rpm for 5 min, resuspended in 2 mL of complete KGM and seeded 50 % each into two new T225 flasks to yield enough cells to then seed 1.5 x 10^6 cells into 9 new T225's. Since continued cell growth for subsequent pull-downs was necessary, one of the 9 T225 flasks was used for another expansion to 2 then 9 T225 flasks. Once cells seeded for pull-downs were ~ 80 % confluent, 4 flasks from each sample were treated for 4 hours with 10 mL of fresh KGM containing 30 µM MG132 proteasome inhibitor in DMSO (MG132, Millipore Sigma., Cat# 474791-5MG or 474791-1MG). The other 4 flasks were each filled with an equal volume of DMSO (vehicle) in 10 mL of KGM. Following treatment, cells were collected as described above and stored in -80°C overnight. A figure outlining the cell culture procedure can be found below (Supplemental Figure 1).

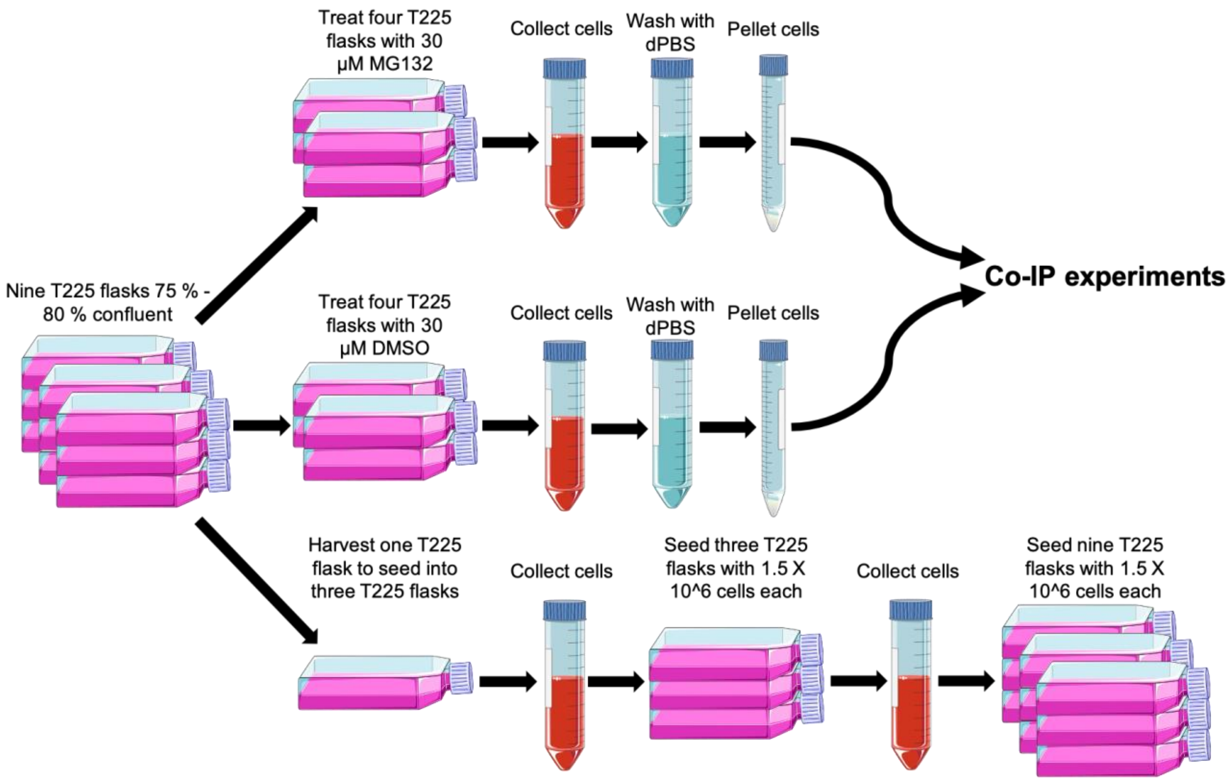


**Supplemental Figure 1** – Generalized process of cell culture for each sample (EPE6, AAE6, PHFK-HA). Once nine flasks were confluent, samples underwent one of three procedures. Top and middle – Treatment of 4 T225 flasks with MG132 and harvesting to collect cells for subsequent Co-IP experiments. Bottom – Trypsinization of a single confluent T225 flask for subsequent expansion into three T225 flasks then once again into nine flasks to continue MG132 and DMSO treatments. Images made using Medical Art by Servier, and free-to-use with attribution (CC BY 3.0).

***Cell Lysis***

To each frozen pellet kept in 15 mL in conical centrifugation tubes (ThermoFisher Sci., Cat# 35296) we added 3 mL of ice-cold lysis buffer consisting of M-PER™ mammalian protein extraction reagent (ThermoFisher Sci., Cat# 78501) supplemented with 1 X HALT$^{TM}$ phosphatase inhibitor cocktail (ThermoFisher Sci., Cat# 78428), 1 X cOmplete EDTA free protease inhibitor (Millipore Sigma., Cat# 4693159001) and 1 mM PMSF and 150 mM NaCl (Fisher Sci., Cat# BP358212). After incubation on a tilting table on ice for 30 min, the lysed cells were separated into two 2 mL low protein binding microcentrifuge tubes (Fisher Sci., Cat# PI88379). Samples were centrifuged twice at 14 000 X g for 15 min at 4°C and thereafter, 5 µL of the resulting supernatant was diluted in 45 µL of ultrapure dd$H_2O$ for protein quantification using the Bradford Assay.

***Western blot***

The following primary antibodies were used: anti-HA 1 : 1000 monoclonal mouse (mAb), (Abcam [HA, C5], Cat# ab181818), anti-E6AP 1 : 1000 polyclonal rabbit (pAb) (Santa Cruz

Antibodies [H-182], Dallas TX, Cat# SC-25509), anti-p53 1 : 500 to 1000 monoclonal rabbit mAb (Abcam [E26], Cambridge UK, Cat# ab32389), anti-HA 1 : 1000 rat mAb conjugated to horseradish peroxidase (HRP) (Roche [3F10], Cat# 1201381900), as well as secondary antibodies: anti-mouse conjugated to HRP 1 : 2000 (Jackson ImmunoResearch Laboratories Inc., West Grove, PA, Cat# 115-035-062) and anti-rabbit conjugated to HRP 1 : 1000 (ThermoFisher Sci., Cat# SA1-200, 40 μg/mL).

All reduced samples (in 1 X SDS loading dye) were run on a mini 4-20% gradient gel (Bio-Rad, Cat# 4561094) for 1 hour and 15 min at a constant voltage of 120 V. Fifteen min prior to the completion of electrophoresis, a polyvinyl difluoride membrane (PVDF, Fisher Sci., Cat# PI88518) was cut to similar dimensions as the mini gel and incubated in methanol for 15 min. Gel was then carefully removed from the casing and placed in ice cold 1 X transfer buffer (100 mL 10 X transfer buffer containing 14.41 g glycine, 3.03 g Tris base, 200 mL methanol) (Fisher Sci., Cat# A454-4) and 700 $dH_2O$. At the same time, the PVDF membrane was transferred into ice-cold 1 X transfer buffer in a separate container and both the gel and membrane were incubated for 10 min. After incubation in transfer buffer, the gel and membrane were placed together in between blotting paper to assemble a transfer sandwich and inserted into an ice-cold tank containing 1 X transfer buffer continuously mixed with a magnetic stir bar. Protein transfer to the membrane lasted for 1 hour at a constant voltage of 100 V and maximum amperage of 0.35 A.

After the transfer, the membrane was removed from the sandwich and incubated for 10 min with 0.05% TBS at room temperature. During the incubation, blocking buffer was prepared (5% (w/v) powdered skim milk (Safeway Pleasanton CA) in 0.05% TBS). Post incubation in 0.05 % TBS-T, the membrane was transferred into a 50 mL conical tube containing 15 mL of prepared blocking buffer. The sample was incubated for 1 hour at room temperature before being transferred into another 50 mL conical cube containing primary antibody in 5 mL of blocking buffer. Once placed in primary antibody, the sample was stored on a rotating rack overnight at 4°C.

The next morning, after antibody removal, samples were washed 3 X 5 min 0.05% TBS-T at room temperature. If the primary antibody did not contain a conjugated HRP tag, the sample was placed in another 50 mL tube containing secondary antibody in 5 mL of blocking buffer for 1 hour at room temperature. After the final incubation in secondary antibody, the samples were washed once again 3 X in 0.05% TBS-T for 5 min. Immediately after the final wash, excess TBS-T was removed by gently placing on a dry paper towel and then placing on plastic wrap. To produce chemiluminescence, samples were incubated in equal volumes of peroxide reagent and luminol/enhancer reagent (Clarity Western ECL Substrate, Bio-Rad, Cat# 1705061) for 1 min. Immediately following incubation, the samples were imaged at 1 X 1 binning for 0.1 seconds, 1, 5, and 20 min intervals using the Ultra-Violet Products Biospectrum Imaging System.

***Co-immunoprecipitation***

For the Co-IP, we intended to use 4 mg of protein extract and 80 µL of Pierce™ Anti-HA Magnetic Beads (ThermoFisher Sci., Cat# 88836) for each condition (i.e. AAE6 pre-treated with MG132). However, the volume of protein lysate required to reach 4 mg total was higher than the volume capacity of the low protein binding microcentrifuge tube. Therefore 2 microcentrifuge tubes, each of them containing 40 µL of beads (pre-washed in supplemented lysis buffer) and 2 mg of protein, we prepared for each condition and incubated on a rotating rack overnight at 4°C. Each sample was then put on a magnetic stand for 4 min at room temperature and the supernatant containing unbound proteins was discarded. Several washes were performed to remove non-specific binders, detergents and protease inhibitors that could interfere with the MS analysis. Beads were resuspended in 1 mL ice-cold, supplemented lysis buffer without inhibitors and incubated for 5-minutes on a rotating rack at 4°C. The samples were incubated 2 min on the magnetic stand at room temperature and the supernatant was discarded. This washing procedure was repeated twice using supplemented lysis buffer without inhibitors and 3 X using 1 mL sterile filtered ice-cold PBS. For each condition, the two tubes were merged together using elution buffer, 100 µL of room temperature 0.2 M glycine (Fisher Sci., Cat# BP3815) pH buffered to 2.5. After a 10 min incubation at room temperature, samples were placed on magnetic stands for 4 min and the eluate was placed into a new 2 mL low protein binding microcentrifuge tube. Immediately after elution, Glycine acidity was neutralized by adding 12 µL of 1 M Tris-HCl (pH 7.5) to each elution sample. Finally, 70% of each sample was stored at -80°C before being analysed by MS facility, while the remaining 30% was used for Western blot.

***Liquid chromatography tandem mass spectrometry (LC-MS/MS) analysis***

Treatment of our transduced keratinocytes at HCMS was as follows. To reduce samples, 2.0 µL of 20 mM Tris(2-carboxyethyl) phosphine hydrochloride solution [pH 1.2-1.5] (Sigma, Cat# 75259-1G) in 50 mM triethylammonium bicarbonate (TEAB) [pH 8.0] (TEAB, Sigma, Cat# T7408) buffer were mixed with 20 µL of sample. Samples were incubated at 37°C for 1 hour and subsequently warmed to room temperature, vortexed and pulse-centrifuged to ensure samples remained at the bottom of each tube. Next, 2.2 µL of freshly prepared 40 mM iodoacetamide (Sigma, Cat# I1149-5G) in 50 mM TEAB was added to each sample for alkylation. Samples with added iodoacetamide were covered with tin foil and incubated at room temperature for 1 hour. After successful reduction and alkylation of samples, trypsin (1:50-1:100 [w/w], Fisher Sci., Cat# SH3023602) was added and incubated at 37°C for 16 hours overnight. After incubation, 1 µL of formic acid (Sigma, Cat# 56302-50ML) was added to each sample and thoroughly vortexed before being placed into a nanoACQUITY ultra performance liquid chromatography (UPLC) system (Waters, Cat# WTF-ARC-2998) to begin LC-MS/MS protein identification. The mass spectrometer used for all LC-MS/MS tests was an Orbitrap Elite™ Hybrid Ion Trap Orbitrap Mass Spectrometer (ThermoFisher Sci., Cat# IQLAAEGAAPFADBMAZQ). The spectra obtained were input into

Proteome Discoverer version 2.4.0.292 for protein identification. The resulting list of proteins was emailed to us as an original “raw” heatmap using Microsoft Excel.
