## Supplementary material for "Virus-host protein-protein interactions between human papillomavirus 16 E6 A1 and D2/D3 sub-lineages: variances and similarities": Dayer et al._Supplemental Table 1

**Supplemental Table 1 – Cellular targets of the human papillomavirus type 16 E6 protein.** The first study encountered in our survey was that by Werness et al. (1990) and the last one that by Oliveira et al. (2018). Most commonly used methods for PPIs were the yeast two hybrid (Y2H) screen, glutathione S-transferase (GST) pulldown, immunoprecipitation (IP) and co-IP. Six papers also reported some form of mass spectrometry (MS) (Nakagawa, Huibregtse 2000, Jeong et al. 2007, Jing et al. 2007, Katzenellenbogen et al. 2007, Vos et al. 2009, White et al. 2012). There is some confusion about protein nomenclature: in the first column (left), we refer to an alias if the UniProt protein nomenclature (The UniProt Consortium 2019) differs from that used in the publication, e.g. in the title and/or abstract; in the other columns, we use the nomenclature adopted by the respective publication. When looking at the binding mechanism, we considered the domain of the targeted protein regarding its amino acid sequence, region (N- or C-terminus) where this information was available. Using the UniProt database (The UniProt Consortium 2019), we have collectively interpreted the data of all 50 binders together in the light of biological functions. Viral processes and immune response, mitogen-activated protein kinases (MAPK)/extracellular signal-regulated kinases (ERK) cascade as well as Wnt (portmanteau of Wingless and int-1, Nusse et al. 1991) and Notch ("notches at the Drosophila wing margin", Artavanis-Tsakonas et al. 1999) signaling pathways—both highly conserved in evolution and embryogenesis-related, were mostly targeted followed by tumour suppressor proteins, DNA damage and repair, apoptosis, adhesion, immortalization and transformation, cell cycle and proliferation, and transcription. In line with these observations, a recent publication dissecting viral associations in patient specimen of the Pan-Cancer Analysis of Whole Genomes Consortium, impaired antiviral defence mechanisms were found to be the driving force for HPV16-related malignancies including cervical, bladder and head and neck cancers (Zapatka et al. 2020).

| Protein Name | Accession Number | Binding Mechanism | Method | Outcome of Interaction | Reference |
|---|---|---|---|---|---|
| (1) Transcriptional adapter 3 (TADA3) alias hADA3 | O75528 | Targets hADA3 to prevent co-activation of p53 | Y2H and GST pulldown | Preventing p53-mediated transactivation of target promoters and p53 stabilization | Kumar et al. 2002 |
| (2) Bcl-2 homologous antagonist/killer (BAK1) | Q16611 | Targets BAK1 through E6AP known to interact with it | GST pulldown | Part of the pro-apoptotic Bcl-2 family; degraded by E6 thereby preventing BAK-induced apoptosis | Thomas and Banks 1999 |
| (3) BRCA1-associated RING domain protein 1 (BARD1) | Q99728 | BARD1 binds to E6 through E6's two zinc finger motifs with zinc finger 1 (AA 30-66) being the most important region for binding | Y2H and IP | Tumour suppressor function through apoptotic signaling inhibited | Yim et al. 2007 |

| Protein Name | Accession Number | Binding Mechanism | Method | Outcome of Interaction | Reference |
|---|---|---|---|---|---|
| (4) Breast cancer type 1 susceptibility protein (BRCA1) | P38398 | Targeted via zinc finger domains of E6 | GST pulldown | Inhibitory telomerase activity inactivated by E6 | Zhang et al. 2005 |
| (5) Golgi-associated PDZ and coiled-coil motif-containing protein (GOPC) alias CAL | Q9HD26 | The PDZ domain of CAL interacts with the PDZ-binding motif of E6; interacts with the E6/E6AP complex enhancing proteasome degradation | GST pulldown and MALDi-TOF | E6 mediates proteasome degradation of CAL through E6AP | Jeong et al. 2007 |
| (6) Histone-arginine methyltransferase CARM (CARM1) | Q86X55 | Not acquired | *In vitro* methyltransferase assay | Prevention of p53-responsive promotors and downregulation of p53 downstream gene expression | Hsu et al. 2012 |
| (7) CREB-binding Protein (CREBBP) alias CBP | Q92793 | E6 binds to 3 regions on CBP and p300: C/H1, C/H3 and C-terminus; binding is independent of p53 | Co-IP | E6 inhibits the activation of p53 and NF[illegible]B by CBP/p300 | Patel et al. 1999 |
| (8) Ubiquitin carboxyl-terminal hydrolase CYLD (CYLD) | Q9NQC7 | Not acquired | EMSA and NF[illegible]B reporter gene assay | E6 mediated ubiquitination and proteasomal degradation of CYLD resulting in hypoxia-induced NF[illegible]B activation | An et al. 2008 |
| (9) Discs large homologue 1 (DLG1) | Q12959 | E6 binds to second PDZ domain via C-terminal XS/TXV/L motif | MBP and GST pulldown | E6 binding to hDLG promotes the transformation of cells | Kiyono et al. 1997 |
| (10) Discs large homologue 4 (DLG4) | P78352 | Binds to E6's C-terminus through its second PDZ motif; last amino acid change from leucin to valine changes affinity | MBP and GST pulldown | Suggested tumour suppressor function; E6 binds induces proteolytic degradation of DLG4 | Handa et al. 2007 |
| (11) Ubiquitin-protein ligase E3A (UBE3A) alias E6AP | Q05086 | Not acquired | Co-IP and GST pulldown | E6 forms a stable complex with E6AP and mediates numerous downstream interactions such as proteolytic degradation of p53 | Huibregtse et al. 1991 |

| Protein Name | Accession Number | Binding Mechanism | Method | Outcome of Interaction | Reference |
|---|---|---|---|---|---|
| (12) Reticulocalbin-2 (RCN2) alias E6BP, ERC55 | Q14257 | E6BP residues 18-29 (VSLEEFLGDY) are the binding site for E6 | Y2H and GST pulldown | Interaction with E6 forms a complex and is thought to be involved with E6 induced transformation and degradation of cellular proteins | Chen et al. 1995, 1998 |
| (13) Signal-induced proliferation-associated 1-like protein 1 (SIPA1L1) alias E6TP1 | O43166 | Binds to E6TP1's C-terminus at residue 194; PDZ domain within E6TP1 has little effect on binding with E6 | Y2H and GST pulldown | Possible tumour suppressor protein; degradation by E6 potentially alters G-associated protein signaling pathways | Gao et al. 1999 |
| (14) FAS-associated death domain protein (FADD) | Q13158 | Targets N-terminus at Serine residue 10, 14, 16 and 18 and Glutamic acid residue 19; site-directed mutants enabled localization of E6-binding to N-terminal end of FADD | Mammalian Y2H and GST pulldown | E6 accelerates depredation of FADD preventing transmission of apoptotic signals through FAS pathway | Filippova et al. 2004 |
| (15) Fibulin-1 (FBLN1) | P23142 | No consensus binding motif identified | Y2H and GST pulldown | Inhibiting Fibulin-1 allowing for invasion and metastasis | Du et al. 2002 |
| (16) E3 ubiquitin-protein ligase HERC2 (HERC2) | O95714 | Interaction with E6 is E6AP dependent | Co-IP and **MS-LC/LC** | Interacts with E6 through formation of complex with E6AP to result in degradation of HERC2 | White et al. 2012 |
| (17) Protein scribble homolog (SCRIB) alias hScrib | Q14160 | Interacts with E6AP in the presence of E6, C-terminus of E6 recognizes PDZ domain of hScrib | GST pulldown and MALDI-MS/MS | Ubiquitination results in degradation and reducing integrity of tight junctions | Nakagawa et al. 2000 |
| (18) Telomerase reverse transcriptase (hTERT) | O94807 | Proximal promotor/regulatory regions (nt position -251 to -88 and +5 to +40) involved with 60% of E6-induced hTERT activity | Telomerase activity and mRNA protection assay | E6 induces increased hTERT activity resulting in maintenance of telomere length | Veldman et al. 2001 |
| (19) Inhibitor of nuclear factor kappa-B kinase subunit beta (IKKB) | O14920 | Not acquired | Co-IP | E6 binding potentially intervenes with NF-kB activation during bacterial or viral infection or DNA damage | Oliveira et al. 2018 |

| Protein Name | Accession Number | Binding Mechanism | Method | Outcome of Interaction | Reference |
|---|---|---|---|---|---|
| (20) Inhibitor of nuclear factor kappa-B kinase subunit epsilon (IKKE) | Q14164 | E6 binds to a location within the first 160 residues of IKKE | Co-IP | E6 binding potentially intervenes with NF-kB activation during TLR9 signaling | Oliveira et al. 2018 |
| (21) Interleukin-1 receptor-associated kinase-like 2 (IRAK2) | O43187 | Not acquired | Co-IP | E6 binding potentially intervenes with NF-kB activation during TLR9 signaling | Oliveira et al. 2018 |
| (22) Interferon regulatory factor 3 (IRF3) alias IRF-3 | Q14653 | IRF-3 residues 109-149 containing ELLG sequence (as in E6AP) | Y2H and GST pulldown | Transcriptional activator; interacts with E6 inhibiting transactivation of IFN-ß | Ronco et al. 1998 |
| (23) Membrane-associated guanylate kinase, WW and PDZ domain-containing protein 1 (MAGI1) alias MAGI-1 | Q96QZ7 | E6 PBM interacts with PDZ1 of MAGI-1 | GST pulldown | Functions in signal transduction and likely a tumour suppressor protein; E6 targets MAGI-1 for proteasomal degradation | Glaunsinger et al. 2000 |
| (24) (MAGI2) alias MAGI-2 | Q86UL8 | PDZ1 domain interacts with E6 most likely through the PDM | *In vitro* degradation assay | Functions in signal transduction; E6 targets MAGI-2 for proteasomal degradation | Thomas et al. 2002 |
| (25) (MAGI3) alias MAGI-3 | Q5TCQ9 | PDZ1 domain interacts with E6 most likely through the PBM | *In vitro* degradation assay | Functions in signal transduction; E6 targets MAGI-3 for proteasomal degradation | Thomas et al. 2002 |
| (26) DNA replication licensing factor MCM7 (MCM7) alias hMCM7 | P33993 | Deletion analysis found E6's N-terminal residues 1-91 bind to hMCM7 C-terminal residues 572-719 | Y2H | Component of replication licensing factors; E6 potentially interferes with its ability to associate with chromatin avoiding G1-phase arrest point | Kukimoto et al. 1998 |
| (27) Methylated-DNA--protein-cysteine methyltransferase (MGMT) | P16455 | MGMT interacts with E6AP through L2G box sequence LLGXXXS/T; PDZ domain present shows potential binding with E6 | GST pull-down and IP | DNA repair protein that protects against mutations; E6 promotes ubiquitination-dependent degradation | Srivenugopal et al. 2002 |
| (28) Myc proto-oncogene protein (MYC) | P01106 | Binds to E6 in an E6AP dependent manner along with E2F1 | GST pulldown | Cellular regulatory processes; E6 binding reduces MYC's half-life and accelerates its degradation | Gross-Mesilaty et al. 1998 |

| Protein Name | Accession Number | Binding Mechanism | Method | Outcome of Interaction | Reference |
|---|---|---|---|---|---|
| (29) Myeloid differentiation primary response protein (MyD88) | Q99836 | Not acquired | Co-IP | E6 binding potentially prevents innate immune receptor signaling | Oliveira et al. 2018 |
| (30) Nuclear factor kappa B subunit 1/2 (NFKB1/2) alias NF[illegible]B | P19838/Q00653 | Not acquired | Immortalization and dual luciferase assays | E6 increased NF-kB levels for baseline and TNF-α by 2- to 3-fold | Vandermark et al. 2012 |
| (31a) Transcriptional repressor (NFX1) alias NFX1-91 | Q12986-3 (NFX1-isoform 3) | NFX1-91 is destabilized by the E6/E6AP complex at NFX-91's C-terminus | Y2H, co-IP and RT-qPCR | Transcriptional repressor of hTERT promoter; E6/E6AP complex destabilizes NFX1-91 through ubiquitination | Gewin et al. 2004 |
| (31b) Transcriptional repressor (NFX1) alias NFX1-91 | Q12986-1 (NFX1-isoform 1) | NFX1-123 is stabilized in the presence of E6 | Y2H, co-IP, GST pulldown and LC-MS/MS | Transcriptional activator of hTERT promoter; E6 may bring NFX1-123 to the hTERT promoter allowing for increased hTERT activation and overexpression | Katzenellenbogen et al. 2007 |
| (32) InaD-like protein (PATJ) | Q8NI35 | E6 binds to the PDZ domain of PATJ (ETQL) | Y2H and co-IP | E6 binds to and targets PATJ for degradation independently of E6AP preventing the formation of a TJ-associated complex Par6-aPKC-PAR3 responsible for regulating kinase activity and formation of tight junctions in polarized cells | Storrs and Silverstein 2007 |
| (33) Paxillin (PAXI) | P49023 | E6 effect likely occurs downstream of paxillin tyrosine phosphorylation and is sensitive to status of actin polymerization | GST pulldown | Transduces signals from plasma membrane to focal adhesions and actin cytoskeleton; E6 disrupts paxillin-mediated actin formation | Tong et al. 1997 |
| (34) E3 ubiquitin-protein ligase PDZRN3 (PDZRN3) | Q9UPQ7 | PDZRN3 interacts with E6 within the PBM | Y2H | When interacting with E6, PDZRN3 is targeted for degradation increasing STAT5-ß activation | Thomas and Banks 2015 |

| Protein Name | Accession Number | Binding Mechanism | Method | Outcome of Interaction | Reference |
|---|---|---|---|---|---|
| (35) Serine/ threonine-protein kinase N1 (PKN1) | Q16512 | E6 binds to C-terminal region of PKN | Y2H and GST pulldown | PKN1 phosphorylates E6, which may allow for E6 to influence Rho-mediated signaling | Gao et al. 2000 |
| (36) Protein arginine N-methyltransferase 1 (PRMT1) | Q99873 | Not acquired | *In vitro* methyltransferase assay | E6 reduces PRMT1-induced methylation of histone H4 at R3 resulting in reduced p53 transactivation | Hsu et al. 2012 |
| (37) Caspase-8 (CASP8) alias Procaspase-8 | Q14790 | Not acquired but both E6 full-length and truncated E6* can bind to procaspase 8 | Mammalian Y2H, GST pulldown, IP and co-IP | E6 targets procaspase 8 for degradation decreasing its interaction with FADD and procaspase 8 dimerization; truncated E6 stabilizes procaspase 8 | Filippova et al. 2007 |
| (38) Tyrosine-protein phosphatase non-receptor type 3 (PTPN3) | P26045 | Interaction between C-terminus of E6 and the PDZ domain of PTPN3 | GST pulldown and LC-MS/MS | Membrane-associated phosphatase; degraded by E6 which prevents tyrosine phosphorylation of growth factor receptors | Jing et al. 2007 |
| (39) Cellular tumour antigen p53 (P53) | P04637 | Not acquired | IP | E6 binds to and degrades p53 | Werness et al. 1990 |
| (40) Histone acetyltransferase p300 (EP300) | Q09472 | Binding domain on E6 between residues 100-147; binds to 3 regions on CBP/p300: C/H1, C/H3 and C-terminus | Co-IP | Coactivator important for cell differentiation and cell cycle progression; E6 prevents the activation of p53 and NFκB via CBP/p300 | Patel et al. 1999 |
| (41) Histone-lysine N-methyltransferase SETD7 (SETD7) | Q8WTS6 | Not acquired | *In vitro* methyltransferase assay | Methylation of histones and non-histone substrates such as p53; inhibition by E6 results in the decrease of p53 stability and activity | Hsu et al. 2012 |
| (42) Telomerase reverse transcriptase (TERT) | O14746 | Not acquired | Modified TRAP | E6 causes ubiquitin-mediated degradation of a telomerase repressing protein | Klingelhutz et al. 1996 |

| Protein Name | Accession Number | Binding Mechanism | Method | Outcome of Interaction | Reference |
|---|---|---|---|---|---|
| (43) Tax-1 binding protein 3 (TX1B3) alias TIP-1 | O14907 | TIP-1 interacts with E6 by binding to the PDZ binding region at E6's terminus | Y2H confirmed with co-IP | TIP-1 interacts with E6 but rather than being degraded, it results in increased activation of RhoA kinase | Hampson et al. 2004 |
| (44) Histone acetyltransferase KAT5 (KAT5) alias TIP60 | Q92993 | Charged residues of the N-terminus of E6 | GST pulldown | E6 destabilizes and degrades TIP60 promoting cell proliferation and cell survival | Jha et al. 2010 |
| (45) Tumor necrosis factor receptor superfamily member 1A (TNR1A) | P19438 | E6 binds to the C-terminal cytoplasmic tail of TNF R1 | IP and mammalian Y2H | E6 inhibits TNF induced apoptosis and formation of the death induced signaling complex (DISC | Filippova et al. 2002 |
| (46) TIR domain-containing adapter molecule 1 (TCAM1) alias TRIF | Q8IUC6 | Not acquired | Co-IP | E6 binding potentially inhibits innate immune functions and antiviral responses | Oliveira et al. 2018 |
| (47) TNF receptor-associated factor 6 (TRAF6) | Q9Y4K3 | Not acquired | Co-IP | E6 binding potentially de-regulates DNA damage response and host immunity | Oliveira et al. 2018 |
| (48a) Tuberin (TSC2) | P49815 | Residues 1-175 and 1251-1807 of TSC2 are required for binding to residues of 260-316 and 428-500 of E6AP | GST pulldown and co-IP | E6 binds to E6AP/TSC2 complex and targets TSC2 for degradation | Zheng et al. 2008 |
| (48b) Tuberin (TSC2) | P49815 | DILG and ELVG domains of Tuberin bind to E6 residues 78-104 | Y2H and GST pulldown | E6 binding causes its ubiquitin-mediated degradation | Lu et al. 2004 |
| (49) Ubiquitin carboxyl-terminal hydrolase 15 (UBP15) alias USP15 | Q9Y4E8 | Not acquired | Targeted MS | Results in increased stability and increased E6 half-life | Vos et al. 2009 |
| (50) DNA repair protein XRCC1 (XRCC1) | P18887 | E6 interacts with the N-terminus of XRCC1 (residues 107-170) | Y2H and co-IP | E6 inhibition prevents ability to maintain genetic integrity and utilize DNA strand break repair mechanisms | Iftner et al. 2002 |
