## Supplementary material for "Virus-host protein-protein interactions between human papillomavirus 16 E6 A1 and D2/D3 sub-lineages: variances and similarities": Dayer et al._Supplemental Table 7-9

**Table 7 – Reactome analysis of the AAE6 interactors selected with the Protein-Pathway method. The pathways with a significant FDR (<0.05) are bolded**

| Pathway name | Entities found | Entities p-value | Entities FDR | UniProt number | Entities protein name |
|---|---|---|---|---|---|
| **(1) Loss of Function of FBXW7 in Cancer and NOTCH1 Signaling** | 3/6 | 3.31E-05 | 0.012 | P62877<br>Q969H0-1<br>Q969H0-4 | E3 ubiquitin-protein ligase *RBX1*<br>F-box/WD repeat-containing protein 7 *FBXW7* isoform 1<br>F-box/WD repeat-containing protein 7 *FBXW7* isoform 4 |
| **(2) FBXW7 Mutants and NOTCH1 in Cancer** | 3/6 | 3.31E-05 | 0.012 | P62877<br>Q969H0-1<br>Q969H0-4 | E3 ubiquitin-protein ligase *RBX1*<br>F-box/WD repeat-containing protein 7 *FBXW7* isoform 1<br>F-box/WD repeat-containing protein 7 *FBXW7* isoform 4 |
| **(3) NOTCH1 Intracellular Domain Regulates Transcription** | 5/48 | 1.30E-04 | 0.019 | P49336<br>P62877<br>Q969H0-1<br>Q969H0-4<br>Q92830 | Cyclin-dependent kinase 8 *CDK8*<br>E3 ubiquitin-protein ligase *RBX1*<br>F-box/WD repeat-containing protein 7 *FBXW7* isoform 1<br>F-box/WD repeat-containing protein 7 *FBXW7* isoform 4<br>Histone acetyltransferase *KAT2A* |
| **(4) Regulation of TP53 Expression** | 2/2 | 1.94E-04 | 0.019 | P04637<br>O75626 | Cellular tumour antigen *P53*<br>PR domain zinc finger protein 1 *PRDM1* |
| **(5) Defective Base Excision Repair Associated with MUTYH** | 2/2 | 1.94E-04 | 0.019 | Q9UIF7-3<br>Q9UIF7-6 | Adenine DNA glycosylase *MUTYH* isoform 3<br>Adenine DNA glycosylase *MUTYH* isoform 6 |
| **(6) Pyrophosphate hydrolysis** | 2/2 | 1.94E-04 | 0.019 | Q9H2U2<br>Q15181 | Inorganic pyrophosphatase 2, mitochondrial *IPYR2*<br>Inorganic pyrophosphatase *IPYR* |
| **(7) Defective MUTYH substrate processing** | 2/2 | 1.94E-04 | 0.019 | Q9UIF7-3<br>Q9UIF7-6 | Adenine DNA glycosylase *MUTYH* isoform 3<br>Adenine DNA glycosylase *MUTYH* isoform 6 |
| **(8) TP53 Regulates Metabolic Genes** | 6/88 | 2.70E-04 | 0.019 | P42345<br>Q9UGJ0<br>P04637<br>P14854<br>P10599<br>Q9HCJ0 | Serine/threonine-protein kinase mTOR *MTOR*<br>5'-AMP-activated protein kinase subunit gamma-2 *AAKG2*<br>Cellular tumour antigen *P53*<br>Cytochrome c oxidase subunit 6B1 *CX6B1*<br>Thioredoxin *THIO*<br>Trinucleotide repeat-containing gene 6C protein *TNR6C* |
| **(9) Constitutive Signaling by NOTCH1 HD+PEST Domain Mutants** | 5/59 | 3.34E-04 | 0.019 | P49336<br>P62877<br>Q969H0-1<br>Q969H0-4<br>Q92830 | Cyclin-dependent kinase 8 *CDK8*<br>E3 ubiquitin-protein ligase *RBX1*<br>F-box/WD repeat-containing protein 7 *FBXW7* isoform 1<br>F-box/WD repeat-containing protein 7 *FBXW7* isoform 4<br>Histone acetyltransferase *KAT2A* |

| Pathway name | Entities found | Entities p-value | Entities FDR | UniProt number | Entities protein name |
|---|---|---|---|---|---|
| **(10) Signaling by NOTCH1 HD+PEST Domain Mutants in Cancer** | 5/59 | 3.34E-04 | 0.019 | P49336<br>P62877<br>Q969H0-1<br>Q969H0-4<br>Q92830 | Cyclin-dependent kinase 8 *CDK8*<br>E3 ubiquitin-protein ligase *RBX1*<br>F-box/WD repeat-containing protein 7 *FBXW7* isoform 1<br>F-box/WD repeat-containing protein 7 *FBXW7* isoform 4<br>Histone acetyltransferase *KAT2A* |
| **(11) Signaling by NOTCH1 PEST Domain Mutants in Cancer** | 5/59 | 3.34E-04 | 0.019 | P49336<br>P62877<br>Q969H0-1<br>Q969H0-4<br>Q92830 | Cyclin-dependent kinase 8 *CDK8*<br>E3 ubiquitin-protein ligase *RBX1*<br>F-box/WD repeat-containing protein 7 *FBXW7* isoform 1<br>F-box/WD repeat-containing protein 7 *FBXW7* isoform 4<br>Histone acetyltransferase *KAT2A* |
| **(12) Constitutive Signaling by NOTCH1 PEST Domain Mutants** | 5/59 | 3.34E-04 | 0.019 | P49336<br>P62877<br>Q969H0-1<br>Q969H0-4<br>Q92830 | Cyclin-dependent kinase 8 *CDK8*<br>E3 ubiquitin-protein ligase *RBX1*<br>F-box/WD repeat-containing protein 7 *FBXW7* isoform 1<br>F-box/WD repeat-containing protein 7 *FBXW7* isoform 4<br>Histone acetyltransferase *KAT2A* |
| **(13) Signaling by NOTCH1 in Cancer** | 5/59 | 3.34E-04 | 0.019 | P49336<br>P62877<br>Q969H0-1<br>Q969H0-4<br>Q92830 | Cyclin-dependent kinase 8 *CDK8*<br>E3 ubiquitin-protein ligase *RBX1*<br>F-box/WD repeat-containing protein 7 *FBXW7* isoform 1<br>F-box/WD repeat-containing protein 7 *FBXW7* isoform 4<br>Histone acetyltransferase *KAT2A* |
| **(14) Signaling by WNT** | 10/299 | 8.48E-04 | 0.044 | P25054<br>Q13237<br>P09497<br>P62877<br>P56545<br>Q9P219<br>P35913<br>Q8N474<br>O14641<br>Q9HCJ0 | Adenomatous polyposis coli protein *APC*<br>cGMP-dependent protein kinase 2 *KGP2*<br>Clathrin light chain B *CLCB*<br>E3 ubiquitin-protein ligase *RBX1*<br>PC-terminal-binding protein 2 *CTBP2*<br>Protein Daple *DAPLE*<br>Rod cGMP-specific 3',5'-cyclic phosphodiesterase subunit beta *PDE6B*<br>Secreted frizzled-related protein 1 *SFRP1*<br>Segment polarity protein dishevelled homolog *DVL2*<br>Trinucleotide repeat-containing gene 6C protein *TNR6C* |

| Pathway name | Entities found | Entities p-value | Entities FDR | UniProt number | Entities protein name |
|---|---|---|---|---|---|
| **(15) Signaling by NOTCH1** | 5/74 | 9.23E-04 | 0.045 | P49336<br>P62877<br>Q969H0-1<br>Q969H0-4<br>Q92830 | Cyclin-dependent kinase 8 *CDK8*<br>E3 ubiquitin-protein ligase *RBX1*<br>F-box/WD repeat-containing protein 7 *FBXW7* isoform 1<br>F-box/WD repeat-containing protein 7 *FBXW7* isoform 4<br>Histone acetyltransferase *KAT2A* |
| (16) Signaling by NOTCH | 8/205 | 0.0011 | 0.049 | P04637<br>P49336<br>P62877<br>Q969H0-1<br>Q969H0-4<br>Q92830<br>Q14186<br>Q9HCJ0 | Cellular tumour antigen *P53*<br>Cyclin-dependent kinase 8 *CDK8*<br>E3 ubiquitin-protein ligase *RBX1*<br>F-box/WD repeat-containing protein 7 *FBXW7* isoform 1<br>F-box/WD repeat-containing protein 7 *FBXW7* isoform 4<br>Histone acetyltransferase *KAT2A*<br>Transcription factor Dp-1 *TFDP1*<br>Trinucleotide repeat-containing gene 6C protein *TNR6C* |
| (17) Negative regulation of TCF-dependent signaling by DVL-interacting proteins | 2/5 | 0.0012 | 0.049 | Q9P219<br>O14641 | Protein Daple *DAPLE*<br>Segment polarity protein dishevelled homolog *DVL2* |
| (18) Activation of NOXA and translocation to mitochondria | 2/5 | 0.0012 | 0.049 | P04637<br>Q14186 | Cellular tumour antigen *P53*<br>Transcription factor Dp-1 *TFDP1* |
| (19) Diseases of Base Excision Repair | 2/7 | 0.0023 | 0.087 | Q9UIF7-3<br>Q9UIF7-6 | Adenine DNA glycosylase *MUTYH* isoform 3<br>Adenine DNA glycosylase *MUTYH* isoform 6 |
| (20) Insulin processing | 3/27 | 0.0026 | 0.091 | P16870<br>Q12840<br>P33176 | Carboxypeptidase E *CBPE*<br>Kinesin heavy chain isoform 5A *KIF5A*<br>Kinesin-1 heavy chain *KINH* |
| (21) Oxidative Stress Induced Senescence | 5/94 | 0.0026 | 0.091 | P04637<br>P01100<br>P10599<br>Q14186<br>Q9HCJ0 | Cellular tumour antigen *P53*<br>Proto-oncogene c-Fos *FOS*<br>Thioredoxin *THIO*<br>Transcription factor Dp-1 *TFDP1*<br>Trinucleotide repeat-containing gene 6C protein *TNR6C* |
| (22) Pre-NOTCH Transcription and Translation | 4/62 | 0.0036 | 0.113 | P04637<br>Q92830<br>Q14186<br>Q9HCJ0 | Cellular tumour antigen *P53*<br>Histone acetyltransferase *KAT2A*<br>Transcription factor Dp-1 *TFDP1*<br>Trinucleotide repeat-containing gene 6C protein *TNR6C* |

| Pathway name | Entities found | Entities p-value | Entities FDR | UniProt number | Entities protein name |
|---|---|---|---|---|---|
| (23) Activation of PUMA and translocation to mitochondria | 2/9 | 0.0038 | 0.113 | P04637<br>Q14186 | Cellular tumour antigen *P53*<br>Transcription factor Dp-1 *TFDP1* |
| (24) Transcriptional Regulation by TP53 | 10/367 | 0.0038 | 0.113 | Q9UGJ0<br>Q92851<br>P04637<br>P14854<br>O75626<br>P01100<br>P42345<br>P10599<br>Q14186<br>Q9HCJ0 | 5'-AMP-activated protein kinase subunit gamma-2 *AAKG2*<br>Caspase-10 *CASP10*<br>Cellular tumour antigen *p53*<br>Cytochrome c oxidase subunit 6B1 *CX6B1*<br>PR domain zinc finger protein 1 *PRDM1*<br>Proto-oncogene c-Fos *FOS*<br>Serine/threonine-protein kinase mTOR *MTOR*<br>Thioredoxin *THIO*<br>Transcription factor Dp-1 *TFDP1*<br>Trinucleotide repeat-containing gene 6C protein *TNR6C* |
| (25) FOXO-mediated transcription | 4/66 | 0.0045 | 0.124 | Q8N139<br>Q969P5<br>P23511<br>P10599 | ATP-binding cassette sub-family A member 6 *ABCA6*<br>F-box only protein 32 *FBX32*<br>Nuclear transcription factor Y subunit alpha *NFYA*<br>Thioredoxin *THIO* |

**Table 8** – **Reactome analysis of the EPE6 interactors selected with the Protein-Pathway method**

| Pathway name | Entities found | Entities p-value | Entities FDR | UniProt number | Entities protein name |
|---|---|---|---|---|---|
| (1) Signaling by Hippo | 3/20 | 4.21E-04 | 0.144 | O14641<br>Q9UDY2<br>Q9GZV5 | Segment polarity protein disheveled homolog *DVL-2*<br>Tight junction protein *ZO-2*<br>WW domain-containing transcription regulator protein 1 *WWTR1* |
| (2) Post-chaperonin tubulin folding pathway | 3/23 | 6.31E-04 | 0.144 | Q13509<br>O75347<br>Q9BTW9 | Tubulin beta-3 chain *TUBB3*<br>Tubulin-specific chaperone A *TBCA*<br>Tubulin-specific chaperone D *TBCD* |
| (3) Activated NTRK3 signals through PI3K | 2/6 | 8.81E-04 | 0.144 | P35568<br>P27986 | Insulin receptor substrate 1 *IRS1*<br>Phosphatidylinositol 3-kinase regulatory subunit alpha *PIK3R1* |
| (4) RNA Polymerase III Transcription Initiation from Type 3 Promoter | 3/28 | 0.0011 | 0.144 | Q16533<br>Q5SXM2<br>A6H8Y1 | snRNA-activating protein complex subunit 1 *SNAPC1*<br>snRNA-activating protein complex subunit 4 *SNAPC4*<br>Transcription factor TFIIIB component B'' homolog *BDP1* |
| (5) HDR through Homologous Recombination (HRR) | 4/66 | 0.0013 | 0.144 | Q99708<br>Q07864<br>P49959<br>P12004 | DNA endonuclease *RBBP8*<br>DNA polymerase epsilon catalytic subunit *POLE*<br>Double strand break repair protein *MRE11*<br>Proliferating cell nuclear antigen *PCNA* |
| (6) Erythrocytes take up oxygen and release carbon dioxide | 2/8 | 0.0016 | 0.144 | P69905<br>P68871 | Hemoglobin subunit alpha *HBA1*<br>Hemoglobin subunit beta *HBB* |
| (7) PI3K/AKT activation | 2/9 | 0.0020 | 0.144 | P35568<br>P27986 | Insulin receptor substrate 1 *IRS1*<br>Phosphatidylinositol 3-kinase regulatory subunit alpha *PIK3R1* |
| (8) RNA Polymerase III Transcription Initiation | 3/36 | 0.0023 | 0.144 | Q16533<br>Q5SXM2<br>A6H8Y1 | snRNA-activating protein complex subunit 1 *SNAPC1*<br>snRNA-activating protein complex subunit 4 *SNAPC4*<br>Transcription factor TFIIIB component B'' homolog *BDP1* |
| (9) HDR through MMEJ (alt-NHEJ) | 2/10 | 0.0024 | 0.144 | Q99708<br>P49959 | DNA endonuclease *RBBP8*<br>Double strand break repair protein *MRE11* |
| (10) Signaling by MET | 4/80 | 0.0027 | 0.144 | Q8N307<br>P27986<br>Q6VN20<br>Q96B97 | Mucin-20 *MUC20*<br>Phosphatidylinositol 3-kinase regulatory subunit alpha *PIK3R1*<br>Ran-binding protein 10 *RANBP10*<br>SH3 domain-containing kinase-binding protein 1 *SH3KBP1* |

| **Pathway name** | **Entities found** | **Entities p-value** | **Entities FDR** | **UniProt number** | **Entities protein name** |
|---|---|---|---|---|---|
| (11) Metabolism of proteins | 25/2012 | 0.0028 | 0.144 | O95786<br>P07550<br>P04637<br>Q8IWV2<br>Q13618<br>Q13217<br>Q9Y297<br>P02671<br>Q92696<br>Q6ZVT0<br>P05019<br>P15088<br>Q8N307<br>P23511<br>Q9UBK2<br>Q14435<br>P12004<br>Q9Y6M0<br>P10599<br>Q9P031<br>Q5JRA6<br>Q13509<br>Q9Y4R7<br>O75347<br>Q9BTW9 | Antiviral innate immune response receptor RIG-I *DDX58*<br>Beta-2 adrenergic receptor *ADRB2*<br>Cellular tumour antigen *p53*<br>Contactin-4 *CNTN4*<br>Cullin-3 *CUL3*<br>DnaJ homolog subfamily C member 3 *DNAJC3*<br>F-box/WD repeat-containing protein 1A *BTRC*<br>Fibrinogen alpha chain *FGA*<br>Geranylgeranyl transferase type-2 subunit alpha *RABGGATA*<br>Inactive polyglycylase *TTLL10*<br>Insulin-like growth factor I *IGF1*<br>Mast cell carboxypeptidase A *CPA3*<br>Mucin-20 *MUC20*<br>Nuclear transcription factor Y subunit alpha *NFYA*<br>Peroxisome proliferator-activated receptor gamma coactivator 1-alpha *PPARGC1A*<br>Polypeptide N-acetylgalactosaminyltransferase 3 *GLNT3*<br>Proliferating cell nuclear antigen *PCNA*<br>Testisin *PRSS21*<br>Thioredoxin *THIO*<br>Thyroid transcription factor 1-associated protein 26 *CCDC59*<br>Transport and Golgi organization protein 1 homolog *MIA3*<br>Tubulin beta-3 chain *TUBB3*<br>Tubulin monoglycylase *TTLL3*<br>Tubulin-specific chaperone A *TBCA*<br>Tubulin-specific chaperone D *TBCD* |
| (12) RNA Polymerase III Abortive and Retractive Initiation | 3/41 | 0.0033 | 0.144 | Q16533<br>Q5SXM2<br>A6H8Y1 | snRNA-activating protein complex subunit 1 *SNAPC1*<br>snRNA-activating protein complex subunit 4 *SNAPC4*<br>Transcription factor TFIIIB component B'' homolog *BDP1* |
| (13) RNA Polymerase III Transcription | 3/41 | 0.0033 | 0.144 | Q16533<br>Q5SXM2<br>A6H8Y1 | snRNA-activating protein complex subunit 1 *SNAPC1*<br>snRNA-activating protein complex subunit 4 *SNAPC4*<br>Transcription factor TFIIIB component B'' homolog *BDP1* |
| (14) Erythropoietin activates Phosphoinositide-3-kinase (PI3K) | 2/12 | 0.0034 | 0.144 | P27986<br>P48736 | Phosphatidylinositol 3-kinase regulatory subunit alpha *PIK3R1*<br>Thyroid transcription factor 1-associated protein 26 *CCDC59* |

| Pathway name | Entities found | Entities p-value | Entities FDR | UniProt number | Entities protein name |
|---|---|---|---|---|---|
| (15) MET activates RAS signaling | 2/12 | 0.0034 | 0.144 | Q8N307<br>Q6VN20 | Mucin-20 *MUC20*<br>Ran-binding protein 10 *RANBP10* |
| (16) Erythrocytes take up carbon dioxide and release oxygen | 2/12 | 0.0034 | 0.144 | P69905<br>P68871 | Hemoglobin subunit alpha *HBA1*<br>Hemoglobin subunit beta *HBB* |
| (17) O2/CO2 exchange in erythrocytes | 2/12 | 0.0034 | 0.144 | P69905<br>P68871 | Hemoglobin subunit alpha *HBA1*<br>Hemoglobin subunit beta *HBB* |
| (18) Carboxyterminal post-translational modifications of tubulin | 3/43 | 0.0037 | 0.145 | Q6ZVT0<br>Q13509<br>Q9Y4R7 | Inactive polyglycylase *TTLL10*<br>Tubulin beta-3 chain *TUBB3*<br>Tubulin monoglycylase *TTLL3* |
| (19) DNA Double-Strand Break Repair | 5/148 | 0.0043 | 0.146 | P04637<br>Q99708<br>Q07864<br>P49959<br>P12004 | Cellular tumour antigen *p53*<br>DNA endonuclease *RBBP8*<br>DNA polymerase epsilon catalytic subunit *POLE*<br>Double strand break repair protein *MRE11*<br>Proliferating cell nuclear antigen *PCNA* |
| (20) TP53 regulates transcription of several additional cell death genes whose specific roles in p53-dependent apoptosis remain uncertain | 2/14 | 0.0046 | 0.146 | P04637<br>Q92696 | Cellular tumour antigen *p53*<br>Geranylgeranyl transferase type-2 subunit alpha *RABGGATA* |
| (21) SEMA3A-Plexin repulsion signaling by inhibiting Integrin adhesion | 2/14 | 0.0046 | 0.146 | O75051<br>Q14563 | Plexin-A2 *PLXNA2*<br>Semaphorin-3A *SEMA3* |
| (22) Transcriptional Regulation by TP53 | 8/367 | 0.0047 | 0.146 | P04637<br>P50750<br>P14854<br>Q99708<br>P49959<br>Q92696<br>P12004<br>P10599 | Cellular tumour antigen *p53*<br>Cyclin-dependent kinase 9 *CDK9*<br>Cytochrome c oxidase subunit 6B1 *COX6B1*<br>DNA endonuclease *RBBP8*<br>Double strand break repair protein *MRE11*<br>Geranylgeranyl transferase type-2 subunit alpha *RABGGATA*<br>Proliferating cell nuclear antigen *PCNA*<br>Thioredoxin *THIO* |
| (23) Toll-like Receptor Cascades | 5/156 | 0.0053 | 0.146 | Q9Y297<br>P02671<br>Q9NWZ3<br>Q13233<br>Q9BT09 | F-box/WD repeat-containing protein 1A *BTRC*<br>Fibrinogen alpha chain *FGA*<br>Interleukin-1 receptor-associated kinase 4 *IRAK4*<br>Mitogen-activated protein kinase kinase kinase 1 *MAP3K1*<br>Protein canopy homolog 3 *CNPY3* |

| Pathway name | Entities found | Entities p-value | Entities FDR | UniProt number | Entities protein name |
|---|---|---|---|---|---|
| (24) Signaling by EGFR | 3/51 | 0.0060 | 0.146 | O43184<br>P27986<br>Q96B97 | Disintegrin and metalloproteinase domain-containing protein 12 *ADAM12*<br>Phosphatidylinositol 3-kinase regulatory subunit alpha *PIK3R1*<br>SH3 domain-containing kinase-binding protein 1 *SH3KBP1* |
| (25) CRMPs in Sema3A signaling | 2/16 | 0.0060 | 0.146 | O75051<br>Q14563 | Plexin-A2 *PLXNA2*<br>Semaphorin-3A *SEMA3* |

**Table 9** – **Reactome analysis of the AAE6 and EPE6 overlapping interactors selected with the Protein-Pathway method**

| Pathway name | Entities found | Entities p-value | Entities FDR | UniProt number | Entities protein name |
|---|---|---|---|---|---|
| (1) TP53 Regulates Metabolic Genes | 3/88 | 0.0005 | 0.094 | P04637<br>P14854<br>P10599 | Cellular tumour antigen *P53*<br>Cytochrome c oxidase subunit 6B1 *CX6B1*<br>Thioredoxin *THIO* |
| (2) Late endosomal microautophagy | 2/34 | 0.0016 | 0.094 | Q9H444<br>P68871 | Charged multivesicular body protein 4b CHMP4B<br>Hemoglobin subunit beta HBB |
| (3) Carboxyterminal post-translational modifications of tubulin | 2/43 | 0.0026 | 0.094 | Q6ZVT0<br>Q9Y4R7 | Inactive polyglycylase *TTLL10*<br>Tubulin monoglycylase *TTLL3* |
| (4) Regulation of TP53 Expression | 1/2 | 0.0035 | 0.094 | P04637 | Cellular tumour antigen *P53* |
| (5) FOXO-mediated transcription | 2/66 | 0.0059 | 0.094 | P23511<br>P10599 | Nuclear transcription factor Y subunit alpha *NFYA*<br>Thioredoxin *THIO* |
| (6) Transcriptional activation of cell cycle inhibitor p21 | 1/4 | 0.0070 | 0.094 | P04637 | Cellular tumour antigen *P53* |
| (7) Transcriptional activation of p53 responsive genes | 1/4 | 0.0070 | 0.094 | P04637 | Cellular tumour antigen *P53* |
| (8) Negative regulation of TCF-dependent signaling by DVL-interacting proteins | 1/5 | 0.0087 | 0.094 | O14641 | Segment polarity protein dishevelled homolog *DVL2* |
| (9) Reelin signalling pathway | 1/5 | 0.0087 | 0.094 | Q96B97 | SH3 domain-containing kinase-binding protein 1 *SH3KBP1* |
| (10) Activation of NOXA and translocation to mitochondria | 1/5 | 0.0087 | 0.094 | P04637 | Cellular tumour antigen *P53* |
| (11) Protein repair | 1/6 | 0.0105 | 0.094 | P10599 | Thioredoxin *THIO* |
| (12) Oxidative Stress Induced Senescence | 2/94 | 0.0117 | 0.094 | P04637<br>P10599 | Cellular tumour antigen *P53*<br>Thioredoxin *THIO* |
| (13) RUNX3 regulates CDKN1A transcription | 1/7 | 0.0122 | 0.094 | P04637 | Cellular tumour antigen *P53* |
| (14) Cell Cycle, Mitotic | 4/536 | 0.0129 | 0.094 | P04637<br>Q13352<br>Q9H444<br>Q96GY3 | Cellular tumour antigen P53<br>Centromere protein R CENPR<br>Charged multivesicular body protein 4b *CHMP4B*<br>Protein lin-37 homolog LIN37 |
| (15) WNT mediated activation of DVL | 1/8 | 0.0139 | 0.094 | O14641 | Segment polarity protein dishevelled homolog *DVL2* |

| Pathway name | Entities found | Entities p-value | Entities FDR | UniProt number | Entities protein name |
|---|---|---|---|---|---|
| (16) Erythrocytes take up oxygen and release carbon dioxide | 1/8 | 0.0139 | 0.094 | P68871 | Hemoglobin subunit beta *HBB* |
| (17) Cargo recognition for clathrin-mediated endocytosis | 2/105 | 0.0145 | 0.094 | O14641<br>Q96B97 | Segment polarity protein dishevelled homolog DVL2<br>SH3 domain-containing kinase-binding protein 1 SH3KBP1 |
| (18) Activation of PUMA and translocation to mitochondria | 1/9 | 0.0157 | 0.094 | P04637 | Cellular tumour antigen *P53* |
| (19) PI5P Regulates TP53 Acetylation | 1/9 | 0.0157 | 0.094 | P04637 | Cellular tumour antigen *P53* |
| (20) Metabolism of proteins | 8/2012 | 0.0158 | 0.094 | P04637<br>P02671<br>Q6ZVT0<br>P23511<br>Q9Y6M0<br>P10599<br>Q9P031<br>Q9Y4R7 | Cellular tumour antigen P53<br>Fibrinogen alpha chain FGA<br>Inactive polyglycylase TTLL10<br>Nuclear transcription factor Y subunit alpha NFYA<br>Testisin PRSS21<br>Thioredoxin THIO<br>Thyroid transcription factor 1-associated protein 26 CCDC59<br>Tubulin monoglycylase TTLL3 |
| (21) ATF6 (ATF6-alpha) activates chaperone genes | 1/10 | 0.0174 | 0.094 | P23511 | Nuclear transcription factor Y subunit alpha *NFYA* |
| (22) Regulation of FOXO transcriptional activity by acetylation | 1/10 | 0.0174 | 0.094 | P10599 | Thioredoxin *THIO* |
| (23) TP53 Regulates Transcription of Death Receptors and Ligands | 1/12 | 0.0209 | 0.094 | P04637 | Cellular tumour antigen *P53* |
| (24) TP53 Regulates Transcription of Caspase Activators and Caspases | 1/12 | 0.0209 | 0.094 | P04637 | Cellular tumour antigen *P53* |
| (25) WNT5:FZD7-mediated leishmania damping | 1/12 | 0.0209 | 0.094 | O14641 | Segment polarity protein dishevelled homolog *DVL2* |
